# Rostral and caudal BLA engage distinct circuits in the prelimbic and infralimbic PFC

**DOI:** 10.1101/2021.12.15.472825

**Authors:** Kasra Manoocheri, Adam G. Carter

## Abstract

Connections from the basolateral amygdala (BLA) to medial prefrontal cortex (PFC) regulate memory and emotion and become disrupted in neuropsychiatric disorders. We hypothesized that the diverse roles attributed to interactions between the BLA and PFC reflect multiple circuits nested within a wider network. To assess these circuits, we first used anatomy to show that the rostral BLA (rBLA) and caudal BLA (cBLA) differentially project to prelimbic (PL) and infralimbic (IL) subregions of the PFC, respectively. We then combined *in vivo* silicon probe recordings and optogenetics to show that rBLA primarily engages PL, whereas cBLA mainly influences IL. Using *ex vivo* whole-cell recordings and optogenetics, we then assessed which neuronal subtypes are targeted, showing that rBLA preferentially drives layer 2 (L2) cortico-amygdalar (CA) neurons in PL, whereas cBLA drives layer 5 (L5) pyramidal tract (PT) cells in IL. Lastly, we used soma-tagged optogenetics to explore the local circuits linking superficial and deep layers of PL, showing how rBLA can also impact L5 PT cells. Together, our findings delineate how subregions of the BLA target distinct networks within the PFC to have different influence on prefrontal output.

## INTRODUCTION

Connections from the basolateral amygdala (BLA) to the medial prefrontal cortex (PFC) regulate memory and emotion (Phelps & LeDoux, 2005; Etkin *et al*., 2011). The BLA sends strong excitatory inputs to the PFC, which contact specific cell types to ultimately influence output of the network (Sotres-Bayon *et al*., 2012; Little & Carter, 2013; Cheriyan *et al*., 2016; Burgos-Robles *et al*., 2017). Dysfunction of these circuits contributes to neuropsychiatric disorders, including post-traumatic stress disorder, major depression, and autism (Gilboa *et al*., 2004; Felix-Ortiz *et al*., 2016; McTeague *et al*., 2020). However, most studies treat the BLA and PFC as monolithic entities, whereas each of these structures actually contains multiple subregions that may have distinct connections and functions (McDonald, 1991; Hintiryan *et al*., 2021). Establishing the interactions between different subregions of the BLA and PFC is important for understanding the myriad of behaviors and disorders involving these areas.

The BLA is often subdivided along the rostrocaudal axis into anterior (magnocellular) and posterior (parvocellular) regions (McDonald, 2003; O’Leary *et al*., 2020). Neurons within the rostral portion of BLA (rBLA) and caudal portion of BLA (cBLA) are implicated in aversive and appetitive behaviors, respectively, suggesting they convey different emotional information to the PFC and other brain regions (Kim *et al*., 2016; Kim *et al*., 2017). The PFC is also subdivided along the dorsoventral axis to include the prelimbic (PL) and infralimbic (IL) regions (Van De Werd *et al*., 2010; Van De Werd & Uylings, 2014), which are functionally opposed in behavioral paradigms like threat conditioning (Sierra-Mercado *et al*., 2011). While PL and IL send diverging projections to subregions of the BLA (McDonald *et al*., 1996; Vertes, 2004), little is known about the reciprocal connection from the rBLA and cBLA to subregions of the PFC.

Recent studies suggest that glutamatergic projections from the BLA activate distinct layers and cell types in PL and IL. BLA inputs to PL preferentially target layer 2 (L2) cortico-amygdalar (CA) neurons, which in turn project back to the BLA to constitute a direct, reciprocal circuit (Little & Carter, 2013; McGarry & Carter, 2016). In contrast, BLA inputs to IL activate layer 5 (L5) pyramidal tract (PT) neurons, which exert top-down control of emotion via projections to the periaqueductal gray (PAG) and other brain regions (Cheriyan *et al*., 2016). One possibility is that BLA neurons send branching connections that target different populations of pyramidal cells in PL and IL. However, an alternative hypothesis is that different populations of neurons in rBLA and cBLA make distinct connections in PL and IL, representing parallel networks linking these brain regions.

In addition to these direct connections, the BLA can influence the PFC by evoking polysynaptic excitation and inhibition. Local connections within and across cortical layers are well studied in other agranular cortices that lack layer 4 (L4) (Hooks *et al*., 2011). For example, layer 2/3 (L2/3) neurons project to deeper layers and target L5 pyramidal cells (Anderson *et al*., 2010; Hirai *et al*., 2012), whereas L5 PT neurons predominantly target other neighboring L5 PT neurons (Morishima & Kawaguchi, 2006). Similarly, stereotyped inhibitory connections across cortical layers are mediated by a variety of interneurons (Saffari *et al*., 2016; Tremblay *et al*., 2016). By engaging networks within and across subregions, the BLA could have complex influences on the PFC, but it remains completely unknown how this differs between rBLA and cBLA inputs.

Here we test the hypothesis that rBLA and cBLA engage distinct networks of pyramidal neurons within the PL and IL of the mouse PFC. Using anatomical tracing, we first show that rBLA projects to PL, whereas cBLA projects to IL, indicating parallel but anatomically separate networks. We then combine *in vivo* recordings and optogenetics to show that rBLA primarily drives PL L2, whereas cBLA engages IL L5. Next, we use *ex vivo* recordings to show the cell-type specificity of these inputs, with rBLA preferentially targeting PL L2 CA neurons and cBLA engaging IL L5 PT neurons. Lastly, we use soma-tagged optogenetics to show how PL L2 CA neurons locally influence PL L5 PT cells. Together, our findings illustrate how rBLA and cBLA evoke activity in PL and IL, demonstrating how subregions of the BLA engage distinct cortical networks in the PFC.

## RESULTS

### Distinct projections of rostral and caudal BLA to prelimbic and infralimbic PFC

To assess if subregions of the PFC receive distinct and spatially segregated inputs from the BLA, we first injected retrograde viruses (AAVrg-tdTomato and AAVrg-GFP) into PL and IL in the same animals (**Fig. 1A**). We observed largely separate populations of PL and IL-projecting neurons across the BLA, with a small population of overlapping dual-projection neurons (PL = 583 ± 59 cells, IL = 694 ± 78 cells, dual-projectors = 246 ± 30 cells, n = 3 animals) (**Fig. 1B**). Quantifying the distributions of labeled cells revealed a smooth gradient along the rostral-caudal axis, with PL-projecting neurons biased towards the rostral BLA (rBLA), and IL-projecting neurons shifted towards the caudal BLA (cBLA) (**Fig. 1C & S1**). These results suggest PL and IL receive inputs from distinct neurons largely residing in rBLA and cBLA, respectively.

**Figure 1:**
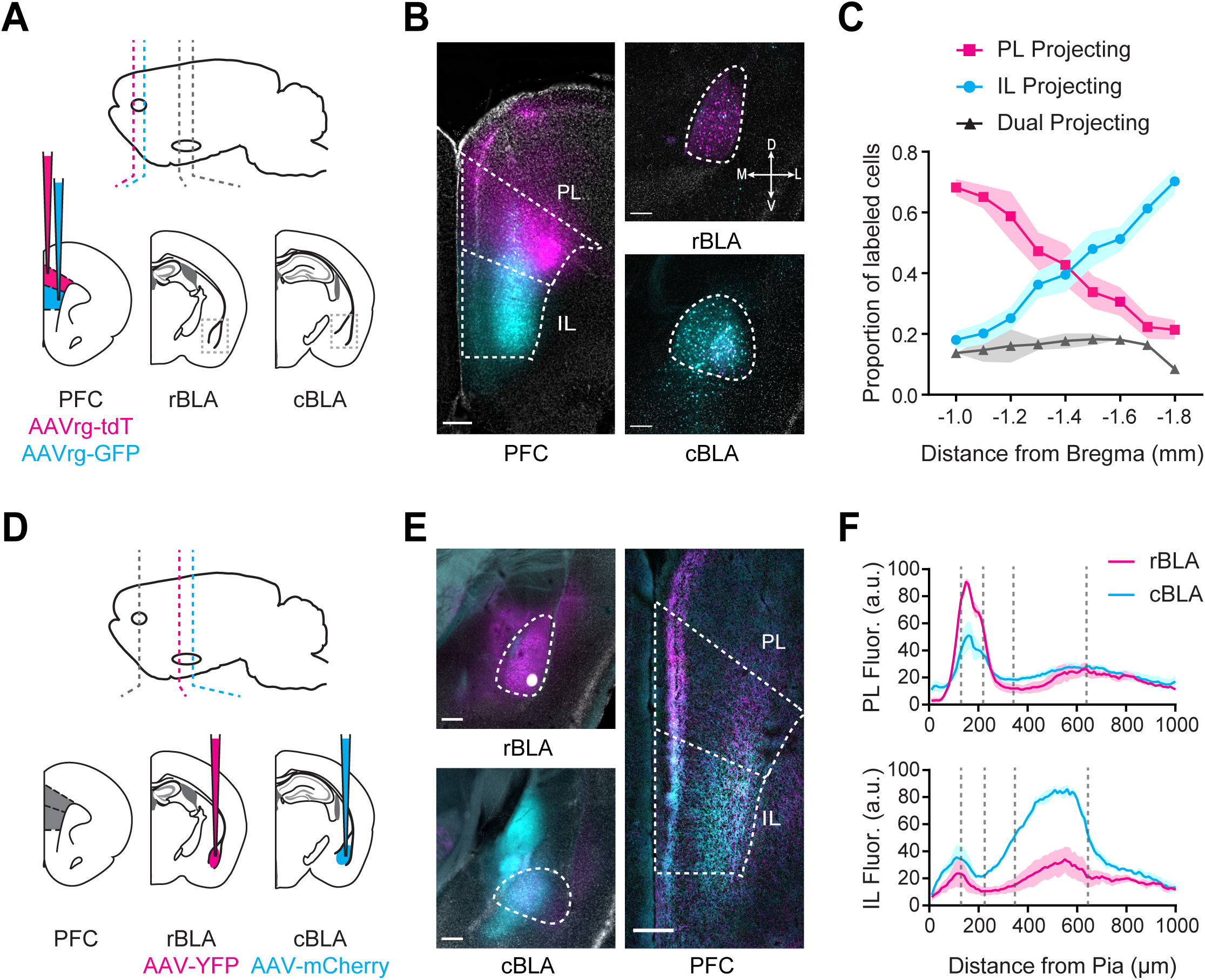
rBLA and cBLA project to different subregions and layers of the PFC. **A)** Schematic for injections of AAVrg-tdTomato into PL (magenta) and AAVrg-GFP into IL (cyan). **B)** *Left,* Image of injection sites in PL (magenta) and IL (cyan). Scale bar = 200 µm. *Right,* Confocal image of rBLA (-1.1 mm from Bregma) and cBLA (-1.7 mm from Bregma). showing retrogradely labeled PL-projecting (magenta) and IL-projecting neurons (cyan). Scale bar = 200 µm. DAPI staining is shown in grey. **C)** Relative distributions of PL-projecting, IL-projecting, and dual-projecting cells in the BLA (n = 3 animals). **D)** Schematic for injections of AAV-YFP into rBLA (magenta) and AAV-mCherry into cBLA (cyan). **E)** *Left,* Images of injection sites in rBLA (magenta) and cBLA (cyan). Scale bar = 250 µm. DAPI staining is shown in grey. *Right,* Confocal image of PL and IL (+2.2 mm from Bregma), showing anterogradely labeled rBLA and cBLA axonal projections. Scale bars = 200 µm. **F)** *Top*, Summary of normalized fluorescence intensity from the pia to white matter of rBLA (magenta) and cBLA (cyan) to PL. *Bottom*, Summary of normalized fluorescence intensity from the pia to white matter of rBLA (magenta) and cBLA (cyan) to IL (n = 3 animals).

To determine how the projections of rBLA and cBLA to the PFC differ, we then injected anterograde viruses (AAV-ChR2-eYFP and AAV-ChR2-mCherry) into the rBLA and cBLA (-1.1 and -1.7 mm from bregma) of the same animals (n = 3) (**Fig. 1D & S1**). Care was taken to avoid injecting virus into the ventral hippocampus, which abuts the cBLA and also projects to the PFC (Liu & Carter, 2018; Liu *et al*., 2020). Consistent with our retrograde labeling, we found rBLA axons primarily in PL and cBLA axons in IL, with distinct laminar targeting (**Fig. 1E**). Quantifying axon fluorescence across layers established that rBLA primarily targeted L2 of PL, whereas cBLA targeted L5 of IL (**Fig. 1F**). Together, these data indicate that rostral and caudal BLA form distinct anatomical connections across layers and the dorsal-ventral axis of the PFC.

### Rostral and caudal BLA evoke distinct activity in prelimbic and infralimbic PFC

To assess differences in the functional impact of rBLA and cBLA inputs on the PFC, we next combined *in vivo* optogenetics and Neuropixels (NP) recordings (Jun *et al*., 2017). We first injected AAV-ChR2-EYFP into either rBLA or cBLA and implanted an optical fiber above the injection site (**Fig. 2A**). After waiting for expression, we acutely recorded light-evoked activity in the PFC of awake, head-fixed mice. The thin dimensions and high density of NP probes allowed us to target L2 or L5 of PL and IL in the same animals (**Fig. 2B**). We confirmed the locations of each NP probe by registering to the Allen common coordinate framework (CCF) (Wang *et al*., 2020) (**Fig. S2**). Our stimulus protocol (5 x 2 ms pulses @ 20 Hz) was similar to previous studies on BLA to PFC connections (Floresco & Tse, 2007; Felix-Ortiz *et al*., 2016). Light-evoked activity was recorded across the dorsal-ventral axis of the PFC and analyzed from NP probes located in either L2 (n = 9 insertions from 6 mice) or L5 (n = 8 insertions from 6 mice) of PL and IL (**Fig. 2C**).

**Figure 2:**
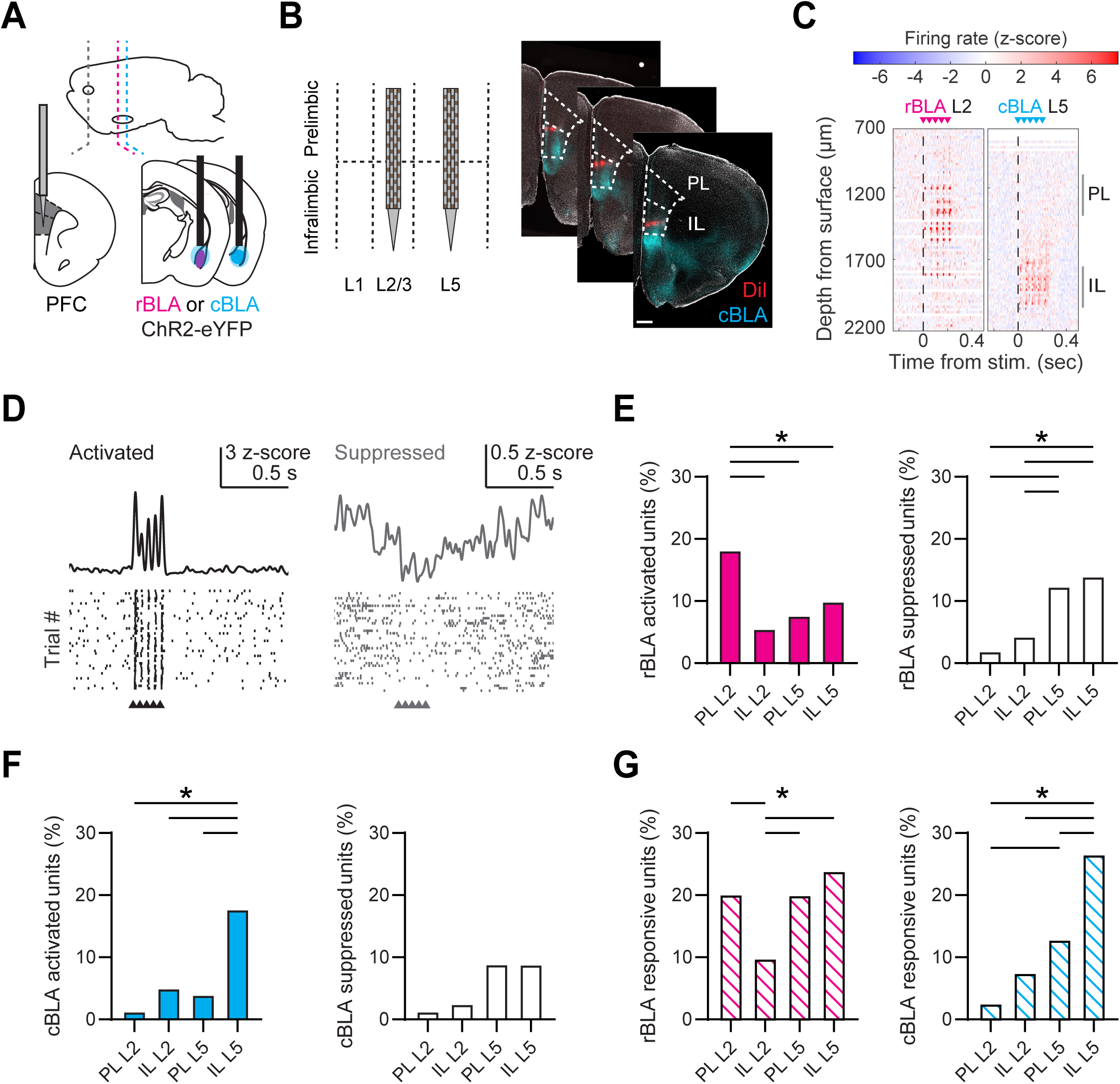
rBLA and cBLA stimulation evoke distinct responses in PL and IL. **A)** Schematic of AAV-ChR2-eYFP injections and fiber optic placement into either rBLA or cBLA, along with Neuropixels (NP) recordings in PFC. **B)** *Left,* Schematic of NP probe insertions into L2/3 and L5 of IL and PL PFC. *Right*, example of NP probe tracts from sequential insertions into L2 and L5 of PFC (red = DiI), along with cBLA axons (cyan = eYFP). Scale bar = 500 µm. DAPI staining is shown in grey. **C)** *Right*, Example rBLA stimulation and NP recording in L2 PFC. Graph shows the average change in spiking activity (z-scored) due to rBLA stimulation across each channel of the NP probe. Arrows denote timing of LED stimulation. *Left*, same but cBLA stimulation and NP recording in L5 PFC. **D)** *Left*, Example single unit with significant BLA-evoked activation, showing average response and raster plot of peri-stimulus activity for 40 trials. *Right*, Similar for a significantly suppressed single unit from BLA stimulation. **E)** *Left,* Percentage of units that were significantly activated by rBLA stimulation in different sub-regions of the PFC. *Right,* Similar for significantly suppressed units. **F)** Similar to (E) for percentages of units affected by cBLA stimulation across sub-regions of the PFC. **G)** Similar to (E) for percentages of units that were responsive (either significantly activated or suppressed) to rBLA stimulation (left) and cBLA stimulation (right). * p < 0.05

To characterize BLA-evoked activity, we used semi-automated spike-sorting to distinguish single units (rBLA-evoked recordings = 979 single units / 1897 total; cBLA-evoked recordings = 596 single units / 1152 total). We found that rBLA and cBLA evoked a variety of responses across the PFC, leading to significantly activated or suppressed units (**Fig. 2D**). The highest proportion of rBLA-activated units were found in PL L2 (PL L2 = 18.2%; IL L2 = 5.5%; PL L5 = 7.9%; IL L5 = 9.9% of single units in subregion), whereas rBLA-suppressed units were enriched in L5 of PL and IL (PL L2 = 1.9%; IL L2 = 4.3%; PL L5 = 12.4%; IL L5 = 14% of single units in subregion) (**Fig. 2E & S3**). In contrast, cBLA-activated units were greatest in IL L5 (PL L2 = 1.3%; IL L2 = 5%; PL L5 = 4%; IL L5 = 17.7% of single units in subregion), with no significant differences in the proportion of cBLA-suppressed units (PL L2 = 1.3%; IL L2 = 2.5%; PL L5 = 8.9%; IL L5 = 8.9% of single units in subregion) (**Fig. 2F & S3**). rBLA stimulation led to more responsive units in PL L2, PL L5, and IL L5 compared to IL L2 (PL L2 = 20.1%; IL L2 = 9.8%; PL L5 = 20.0%; IL L5 = 23.9% of single units in subregion), whereas cBLA stimulation mainly drove responses in IL L5 (PL L2 = 2.6%; IL L2 = 7.5%; PL L5 = 12.9%; IL L5 = 26.6% of single units in subregion) (**Fig. 2G**). We observed no significant differences in latency to spike after each LED pulse in activated units (**Fig. S3**). Together, these findings indicate that rBLA and cBLA drive distinct activity in PL and IL, with rBLA primarily activating PL L2, but also unexpectedly generating mixed activation and suppression across L5, and cBLA predominantly activating IL L5 with fewer responses elsewhere.

### Rostral and caudal BLA target different cell types in prelimbic and infralimbic PFC

Our *in vivo* recordings showed that rBLA and cBLA engage different subregions and layers of PFC, but did not reveal which cell types are targeted. Recent work shows BLA inputs can selectively target L2 cortico-amygdalar (CA) neurons in PL or L5 pyramidal tract (PT) neurons in IL (Little & Carter, 2013; Cheriyan *et al*., 2016; McGarry & Carter, 2016). We next used whole-cell recordings and optogenetics to test whether rBLA and cBLA differentially engaged these cell types in PL and IL. We injected ChR2-expressing virus (AAV-ChR2-EYFP) into either rBLA or cBLA to visualize and stimulate axons in the PFC (**Fig. 3A**) (Little & Carter, 2013; McGarry & Carter, 2016). In the same mice, we also co-injected retrogradely transported, fluorescently-tagged cholera toxin subunit B (CTB) into either rBLA or cBLA and the periaqueductal gray (PAG) to label CA and PT cells, respectively (**Fig. 3A**) (Collins *et al*., 2018; Liu & Carter, 2018). After waiting for expression and transport, we observed CA cell labeling in L2 and PT cell labeling in L5 across PL and IL (**Fig. 3B**). We then used wide-field illumination to activate rBLA or cBLA inputs, and measured synaptic responses at CA and PT cells in TTX (1 µM), 4-AP (100 µM) and high extracellular Ca^2+^ (4 mM) to isolate monosynaptic connections (Petreanu *et al*., 2009). To control for variability in virus expression between slices and animals, we compared the ratio of responses referenced to a specific cell-type (either PL L2 CA for rBLA or IL L5 PT for cBLA) (**Fig. 3B**).

**Figure 3:**
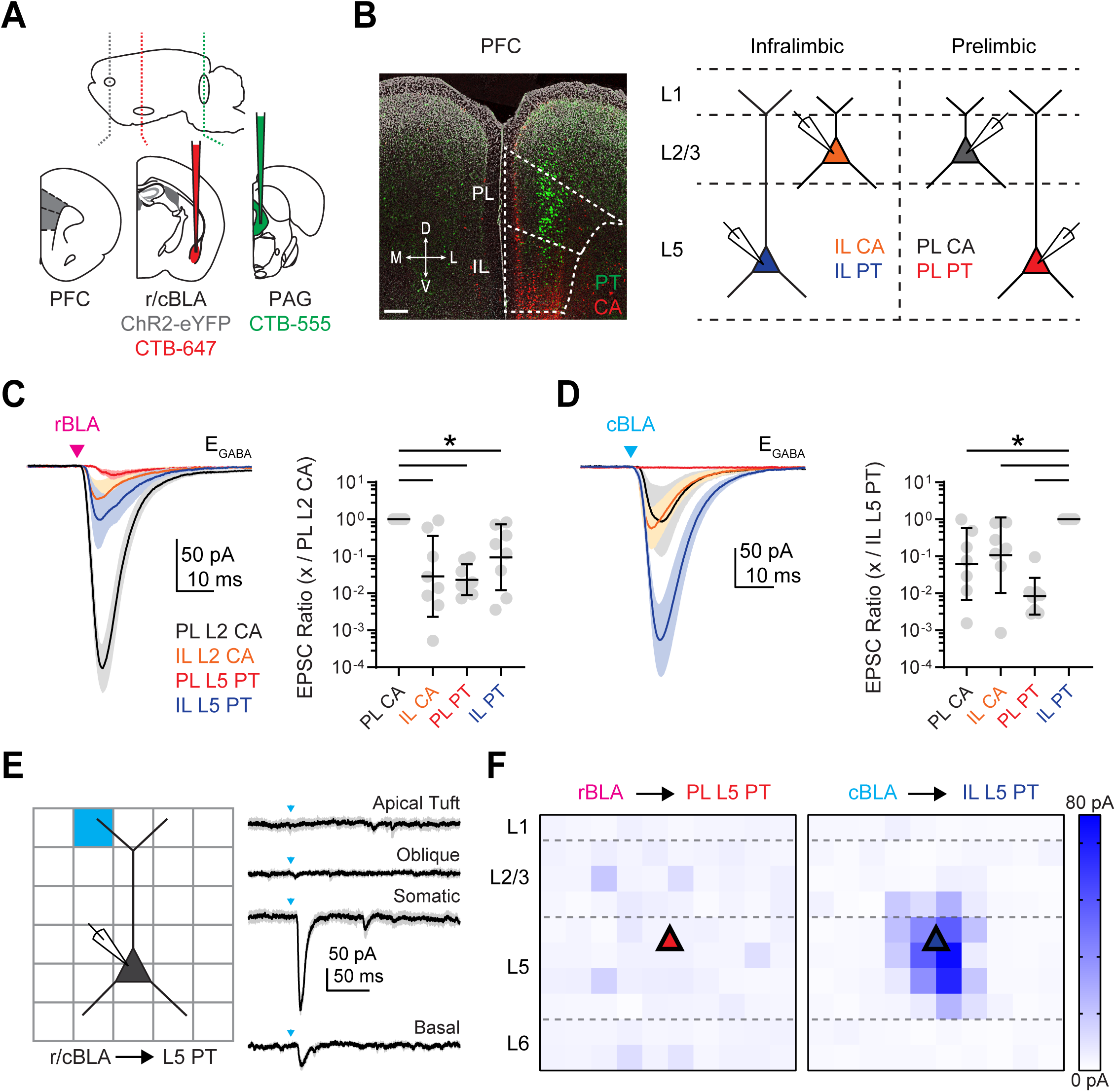
rBLA targets PL L2 CA neurons and cBLA targets IL L5 PT neurons. **A)** *Left,* Schematic for injections of AAV-ChR2-eYFP and CTB-647 into rBLA/cBLA and CTB-555 into PAG. *Right,* Example image of CTB-647 labeled CA neurons (red) and CTB-555 labeled PT neurons (green) in PFC (+2.2 mm from Bregma). Scale bar = 250 µm. DAPI staining is shown in grey. **B)** Experimental scheme for voltage-clamp recordings. PL L2 CA, IL L2 CA, PL L5 PT, and IL L5 PT neurons were recorded in the presence of TTX and 4-AP to isolate monosynaptic connections from rBLA or cBLA. **C)** *Left,* Average rBLA-evoked EPSCs at L2 CA and L5 PT neurons in PL and IL. Magenta arrow = light stimulation. *Right,* Summary of EPSC amplitude ratios, comparing PL L2 CA neurons versus paired IL L2 CA (n = 8), IL L5 PT (n = 8), and PL L5 PT (n = 8) neurons (5 animals). **D)** Similar to (C) for cBLA-evoked EPSCs, including comparison of IL L5 PT neurons with paired IL L2 CA (n = 7), PL L2 CA (n = 7), and PL L5 PT (n = 8) neurons (6 animals). **E)** *Left,* Experimental scheme for sCrACM experiments, showing pseudorandom illumination (blue light) of 75 x 75 µm squares across L1 to L6 of PFC while recording rBLA-evoked EPSCs at PL L5 PT neurons and cBLA-evoked EPSCs and IL L5 PT neurons in the presence of TTX and 4-AP. *Right,* Example cBLA-evoked responses at IL L5 PT neuron, including at apical tuft, oblique, somatic, and basal dendrite locations. **F)** *Left,* Average rBLA-evoked EPSC maps for PL L5 PT neurons (n = 6 cells; 3 animals). *Right,* Average cBLA-evoked EPSC maps for IL L5 PT neurons (n = 6 cells, 4 animals). Triangle denotes typical cell body location. * p < 0.05

Taking this approach, we observed that rBLA-evoked excitatory postsynaptic currents (EPSCs) at -60 mV were much larger at PL L2 CA neurons compared to all other cell types in the different layers and sub-regions (EPSC Ratio: IL L2 CA vs. PL L2 CA = 0.03, geometric standard deviation factor [GSD] = 12.3, p = 0.0003; n = 8 pairs; PL L5 PT vs. PL L2 CA = 0.02, GSD = 2.6, p = 0.0001; n = 8 pairs; IL L5 PT vs. PL L2 CA = 0.09, GSD = 7.7, p = 0.006; n = 8 pairs; 5 animals) (**Fig. 3C & S4**). In contrast, cBLA-evoked EPSCs were much larger at IL L5 PT neurons compared to all other cell types (EPSC Ratio: PL L2 CA vs. IL L5 PT = 0.06, GSD = 9.3, p = 0.005; n = 7 pairs; IL L2 CA vs. IL L5 PT = 0.11, GSD = 10.5, p = 0.014; n = 7 pairs; PL L5 PT vs. IL L5 PT = 0.008, GSD = 3.1, p < 0.0001; n = 8 pairs; 6 animals) (**Fig. 3D & S4**). We also confirmed that rBLA inputs target PL L2 CA neurons over other L2 projection neurons and that cBLA inputs target IL L5 PT over other L5 projection neurons (**Fig. S4**). These findings confirm that sub-regions of BLA preferentially target different layers and cell-types in the PFC, and further indicate that rBLA primarily synapses onto PL L2 CA cells, and cBLA primarily contacts IL L5 PT cells.

Other long-range excitatory inputs to the PFC can evoke unique responses by selectively targeting the apical dendrites of pyramidal cells (Anastasiades *et al*., 2021). Because our anatomy showed prominent BLA axons in superficial layers, we tested the possibility that BLA inputs also contact the apical dendrites of L5 PT cells using sub-cellular Channelrhodopsin-Assisted Circuit Mapping (sCrACM) in TTX and 4-AP (Petreanu *et al*., 2009), stimulating across a grid aligned to the pia and soma (Anastasiades *et al*., 2021) (**Fig. 3E**). We found strong cBLA input to the basal and peri-somatic dendrites of IL L5 PT cells, but no rBLA input to either the apical or basal dendrites of PL L5 PT cells, unlike rBLA inputs to PL L2 CA cells (**Fig. 3F & S4**). These findings indicate that rBLA and cBLA primarily make connections close to the soma of pyramidal cells in the PFC, and do not target the apical dendrites of deep layer cells.

### Rostral and caudal BLA activate different networks in prelimbic and infralimbic PFC

Having established which cell types are targeted by the BLA, we next assessed how they are engaged during more physiological conditions and stimulation patterns. We were particularly interested in how rBLA and cBLA inputs evoked excitation and inhibition at different cell types in the PL and IL, in order to understand the mechanisms behind the *in vivo* responses we measured.

We stimulated rBLA or cBLA inputs with the same repetitive trains as in our *in vivo* recordings (5 pulses of 473 nm light for 2 ms at 20 Hz) and recorded EPSCs at -60 mV (E_GABA_) and IPSCs at +15 mV (E_AMPA_) in the presence of CPP (10 µM) to block NMDA receptors. For rBLA inputs, we recorded pairs of PL L2 CA neurons and either IL L2 CA, PL L5 PT, or IL L5 PT neurons (**Fig. 4A & S5**). We found that rBLA-evoked EPSCs were again largest at PL L2 CA neurons (EPSC Ratio: IL L2 CA vs. PL L2 CA = 0.05, GSD = 3, p < 0.0001, n = 8 pairs; PL L5 PT vs. PL L2 CA = 0.11, GSD = 4.6, p = 0.0002, n = 12 pairs; IL L5 PT vs. PL L2 CA = 0.18, GSD = 3.4, p = 0.004, n = 7 pairs, 10 animals) (**Fig. 4B & C**). In contrast, rBLA-evoked IPSCs were of comparable amplitude at PL L2 CA, PL L5 PT, and IL L5 PT neurons (IPSC Ratio: IL L2 CA vs. PL L2 CA = 0.04, GSD = 7.2, p = 0.004, n = 7 pairs; PL L5 PT vs. PL L2 CA = 1.1, GSD = 2.6, p > 0.99, n = 12 pairs; IL L5 PT vs. PL L2 CA = 0.97, GSD = 2.8, p > 0.99, n = 7 pairs, 10 animals) (**Fig. 4B & C**). For cBLA inputs, we recorded pairs of IL L5 PT neurons and either PL L2 CA, PL L5 PT or IL L2 CA neurons (**Fig. 4D & S5**). We found that both EPSCs and IPSCs were strongly biased onto IL L5 PT neurons compared to all other cell types (EPSC Ratio: PL L2 CA vs. IL L5 PT = 0.06, GSD = 3.7, p = 0.003; n = 7 pairs; IL L2 CA vs. IL L5 PT = 0.08, GSD = 3.4, p = 0.02; n = 6 pairs; PL L5 PT vs. IL L5 PT = 0.005, GSD = 8.7, p < 0.0001; n = 8 pairs, 9 animals) (IPSC Ratio: PL L2 CA vs. IL L5 PT = 0.006, GSD = 6.4, p = 0.0003; n = 7 pairs; IL L2 CA vs. IL L5 PT = 0.002, GSD = 3.6, p < 0.0001; n = 6 pairs; PL L5 PT vs. IL L5 PT = 0.01, GSD = 7.8, p = 0.006; n = 8 pairs, 9 animals) (**Fig. 4E & F**). We also observed differences in excitation / inhibition (E/I) ratios, with rBLA inputs onto PL L2 CA showing a higher E/I ratio than rBLA inputs onto PL L5 PT and IL L5 PT neurons (**Fig. S5**). These results confirm that rBLA excites PL L2 CA neurons, but inhibits multiple cell types, whereas cBLA primarily excites and inhibits IL L5 PT.

**Figure 4:**
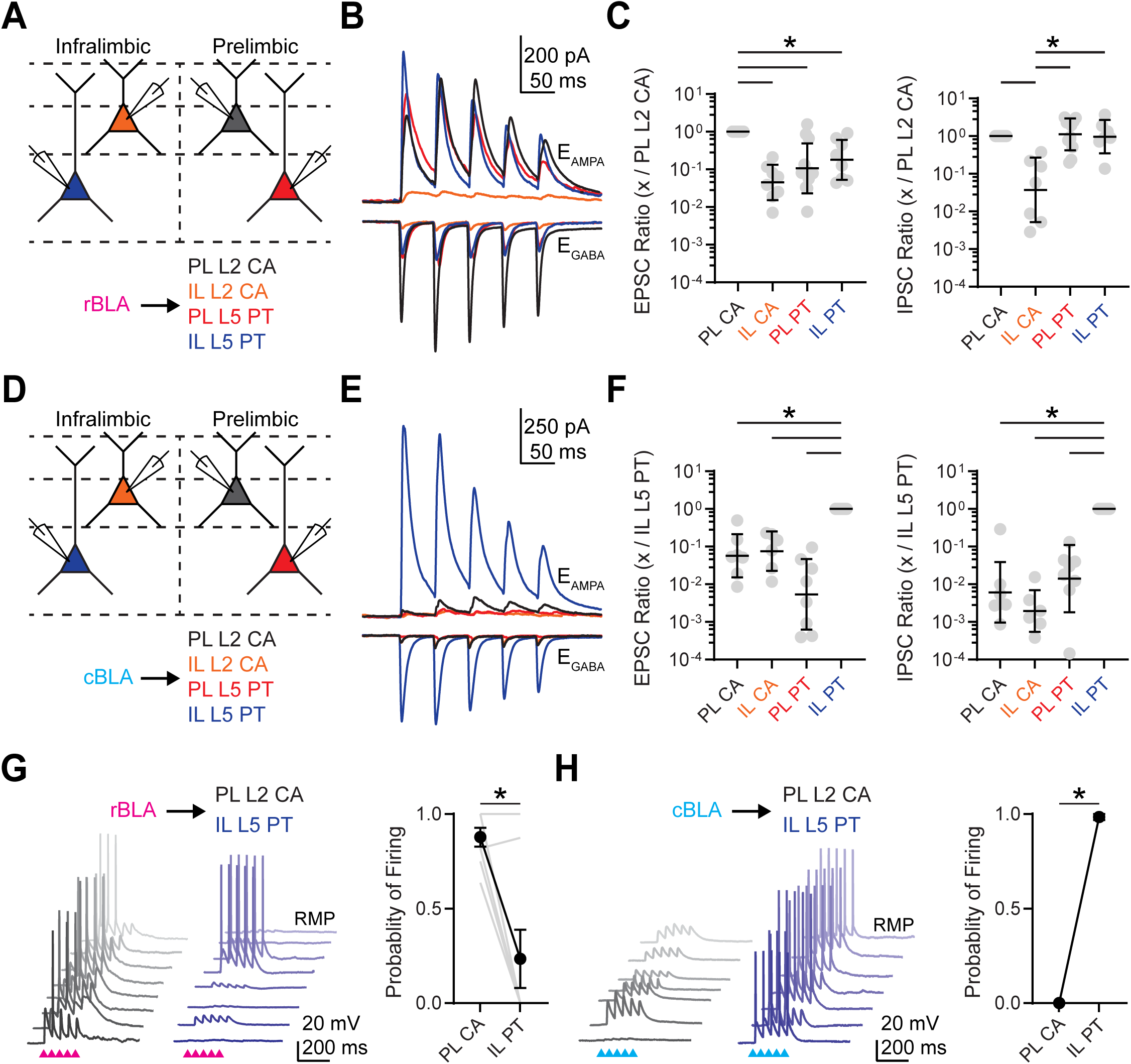
rBLA and cBLA activate distinct projection neurons in PL and IL. **A)** Experimental scheme for recording rBLA-evoked excitation and inhibition from PL L2 CA (black), IL L2 CA (orange), PL L5 PT (red) and IL L5 PT (blue) neurons. **B)** Average rBLA-evoked EPSCs and IPSCs at PL L2 CA (black), IL L2 CA (orange), PL L5 PT (red) and IL L5 PT neurons (blue). PL L2 CA average traces reflect data from all sets of pairs (n = 19 PL L2 CA neurons, 8 IL L2 CA neurons, 12 PL L5 PT neurons, and 8 IL L5 PT neurons, 10 animals). **C)** *Left,* Summary of rBLA-evoked EPSC_1_ amplitude ratios for PL L2 CA, IL L2 CA, PL L5 PT, and IL L5 PT responses divided by PL L2 CA responses. *Right,* Similar for IPSC_1_ amplitude ratios. Grey points denote ratios of individual pairs. **D – F)** Similar to (A – C) for cBLA inputs onto PL L2 CA, IL L2 CA, PL L5 PT, and IL L5 PT neurons divided by IL L5 PT responses (n = 16 IL L5 PT neurons, 7 PL L2 CA neurons, 8 PL L5 PT neurons, and 6 IL L2 CA neurons, 9 animals). **G)** *Left*, Example rBLA-evoked EPSPs and APs from each pair of PL L2 CA and IL L5 PT neurons recorded in current-clamp from RMP. Arrows denote light stimulation. *Right*, Summary of rBLA-evoked AP probability for PL L2 CA and IL L5 PT neurons, where grey lines denote pairs (n = 8 cells, 5 animals). **H)** Similar to (G) for cBLA-evoked EPSPs and APs from pairs of PL L2 CA and IL L5 PT neurons (note that grey lines are omitted) (n = 7 cells, 3 animals). * p < 0.05

We then tested how rBLA and cBLA inputs drive cell-type specific firing of the main recipient cell types in PL and IL. Projection neurons in the PFC have distinct intrinsic properties (Ferreira *et al*., 2015), with CA neurons resting at very hyperpolarized potentials (McGarry & Carter, 2016) and PT cells displaying robust h-current (Dembrow *et al*., 2010; Anastasiades *et al*., 2018). Taking these properties into account, we recorded responses in current-clamp at resting membrane potential (RMP). We found that rBLA inputs evoked significantly more action potential firing of PL L2 CA neurons compared to IL L5 PT neurons in the same slices (Firing probability: PL L2 CA = 0.88 **±** .05, IL L5 PT = 0.23 **±** .15, p = 0.016; n = 8 pairs, 5 animals) (**Fig. 4G**). In contrast, cBLA inputs activated only IL L5 PT neurons and never activated PL L2 CA neurons (Firing probability: PL L2 CA = 0, IL L5 PT = 0.98 **±** .02, p = 0.03; n = 7 pairs, 3 animals) (**Fig. 4H**). Together, these findings indicate rBLA and cBLA engage distinct projection neuron networks in the PFC, supporting our hypothesis that BLA consists of functionally distinct rostral and caudal divisions.

### Rostral BLA evokes disynaptic excitation and inhibition of PL L5 PT neurons

While our *in vivo* recordings indicate that rBLA inputs primarily target PL L2, we also observed a substantial number of activated and suppressed units in PL L5 and IL L5. Similarly, previous studies have found that BLA influences the firing of PL L5 PT cells to shape PFC output (Huang *et al*., 2019). Because we found no monosynaptic connections onto PL L5 PT neurons, we hypothesized a role for local connections. To test this, we combined current-clamp recordings of PL L2 CA neurons with voltage-clamp recordings of PL L5 PT cells (**Fig. 5A**). We first illuminated rBLA inputs to PL L2 with trains that evoked sub-threshold or supra-threshold activity at PL L2 CA neurons. In the same slices, we recorded rBLA-evoked EPSCs at E_GABA_ and IPSCs at E_AMPA_ from PL L5 PT cells. We observed no responses at subthreshold intensities, but robust responses at suprathreshold intensities, suggesting polysynaptic activity from L2 to L5 is involved in rBLA-evoked responses at PL L5 PT cells (EPSC: sub = -9 **±** 4 pA, supra = -113 **±** 24 pA, p = 0.02; n = 8 pairs; IPSC: sub = 17 **±** 5 pA, supra = 508 **±** 101 pA, p = 0.008; n = 8 pairs; 4 animals) (**Fig. 5B & C**). In contrast, we observed no responses from IL L5 PT cells at either subthreshold or suprathreshold intensities, suggesting activity does not spread from PL L2 to IL L5 in the slice, and that rBLA instead projects directly to IL L5 (EPSC: sub = -4 **±** 3 pA, supra = -7 **±** 4 pA, p = 0.055; n = 8 pairs; IPSC: sub = -7 **±** 2 pA, supra = -11 **±** 3 pA, p = 0.64; n = 8 pairs; 3 animals) (**Fig. 5D-F**). In order to confirm that cBLA does not engage PL L5 via IL L5, we also activated cBLA inputs in IL L5, and observed no EPSCs at either sub-threshold or supra-threshold intensities, and only minimal IPSCs at supra-threshold intensities, suggesting polysynaptic activity also does not spread from IL L5 to PL L5 (EPSC: sub = -4 **±** 2 pA, supra = -13 **±** 5 pA, p = 0.055; n = 9 pairs; IPSC: sub = 7 **±** 2 pA, supra = 54 **±** 22 pA, p = 0.027; n = 9 pairs; 3 animals) (**Fig. 5G-I**). These findings suggest that rBLA, but not cBLA, influences PL L5 PT cells via local connections across layers of the PFC.

**Figure 5.**
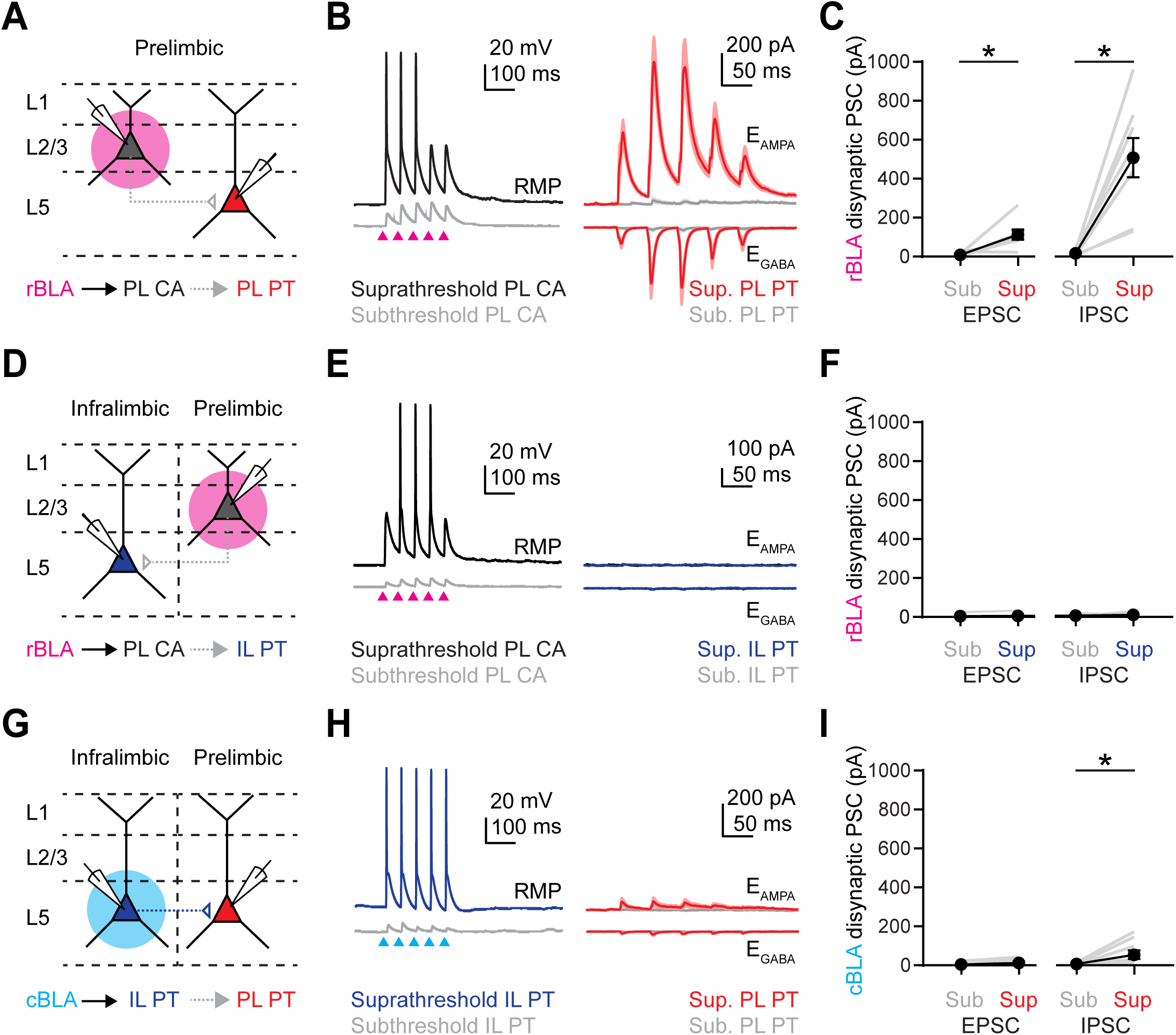
rBLA but not cBLA engages PL L5 PT neurons via local networks. **A)** Experimental scheme for rBLA-evoked responses at PL L2 CA neurons (black) and polysynaptic responses at PL L5 PT neurons (red). **B)** *Left,* Example rBLA-evoked EPSPs and APs from PL L2 CA neurons recorded in current-clamp at RMP with subthreshold (grey) and suprathreshold (black) stimulation. *Right,* Average rBLA-evoked EPSCs and IPSCs from PL L5 PT neurons at the same subthreshold (grey) and suprathreshold (red) stimulation. **C)** Summary of rBLA-evoked EPSC_1_ and IPSC_1_ amplitudes at PL L5 PT neurons. Grey lines denote subthreshold and suprathreshold responses in the same neuron (n = 8 cells, 4 animals). **(D – F)** Similar to (A – C) for rBLA-evoked firing at PL L2 CA neurons (black) and lack of polysynaptic responses at IL L5 PT neurons (blue) (n = 8 cells, 3 animals). **(G – I)** Similar to (A – C) for cBLA-evoked firing at IL L5 PT neurons (blue) and lack of polysynaptic responses at PL L5 PT neurons (red) (n = 9 cells, 3 animals). * p < 0.05

### Layer 2 CA neurons selectively engage Layer 5 PT neurons in PL PFC

We next sought to determine how L2 CA neurons influence L5 PT neurons across layers of the PL PFC and whether they target other projection neurons. We first injected AAVrg-Cre into the BLA, as well as AAV-DIO-ChR2 into the PFC, allowing for expression of ChR2 in CA neurons (**Fig. S6**). We then recorded CA-evoked responses at triplets of pyramidal cells, observing robust EPSCs and IPSCs in L2 and L5 but not L3 (**Fig. S6**) In separate experiments, we also recorded CA-evoked responses at pairs of L5 pyramidal cells, and observed biased EPSCs and IPSCs at PT cells compared to neighboring IT cells (**Fig. S6**). These results suggest that CA neurons can directly engage L5 PT cells, but their interpretation is complicated by the presence of presynaptic CA neurons in both L2 and L5, making it unclear if connections arise from within L5 itself, or are inherited from superficial layers from L2 (Cheriyan *et al*., 2016; Avesar *et al*., 2018).

To overcome this challenge, we next used the soma-targeted opsin st-ChroME (Mardinly *et al*., 2018) to selectively activate L2 CA neurons and record evoked responses at L5 pyramidal cells. We injected AAVrg-Cre into the BLA, as well as AAV-DIO-st-ChroME into the PFC, allowing for expression in CA neurons (**Fig. 6A**). We illuminated presynaptic cells in either L2 or L5 with a bar of light (**Fig. 6B**), demonstrating their layer-specific activation (**Fig. 6C**). We then stimulated L2 CA neurons with trains of light at 20 Hz and recorded evoked responses at from pairs of labeled PL L5 IT or PL L5 PT cells (**Fig. 6D**). We observed robust L2 CA-evoked EPSCs and IPSCs, which were again strongly biased onto L5 PT cells compared to neighboring L5 IT cells (EPSC: PL L5 PT = -67 **±** 15 pA, PL L5 IT = -4 **±** 1 pA, p = 0.0005; n = 12 pairs; 7 animals; IPSC: PL L5 PT = 229 **±** 92 pA, PL L5 IT = 20 **±** 15 pA, p = 0.03; n = 10 pairs; 6 animals) (**Fig. 6E & S6**). These results indicate that activation of PL L2 CA cells, which are the main cell type activated by rBLA input, in turn generates biased excitation and inhibition at PL L5 PT cells.

**Figure 6.**
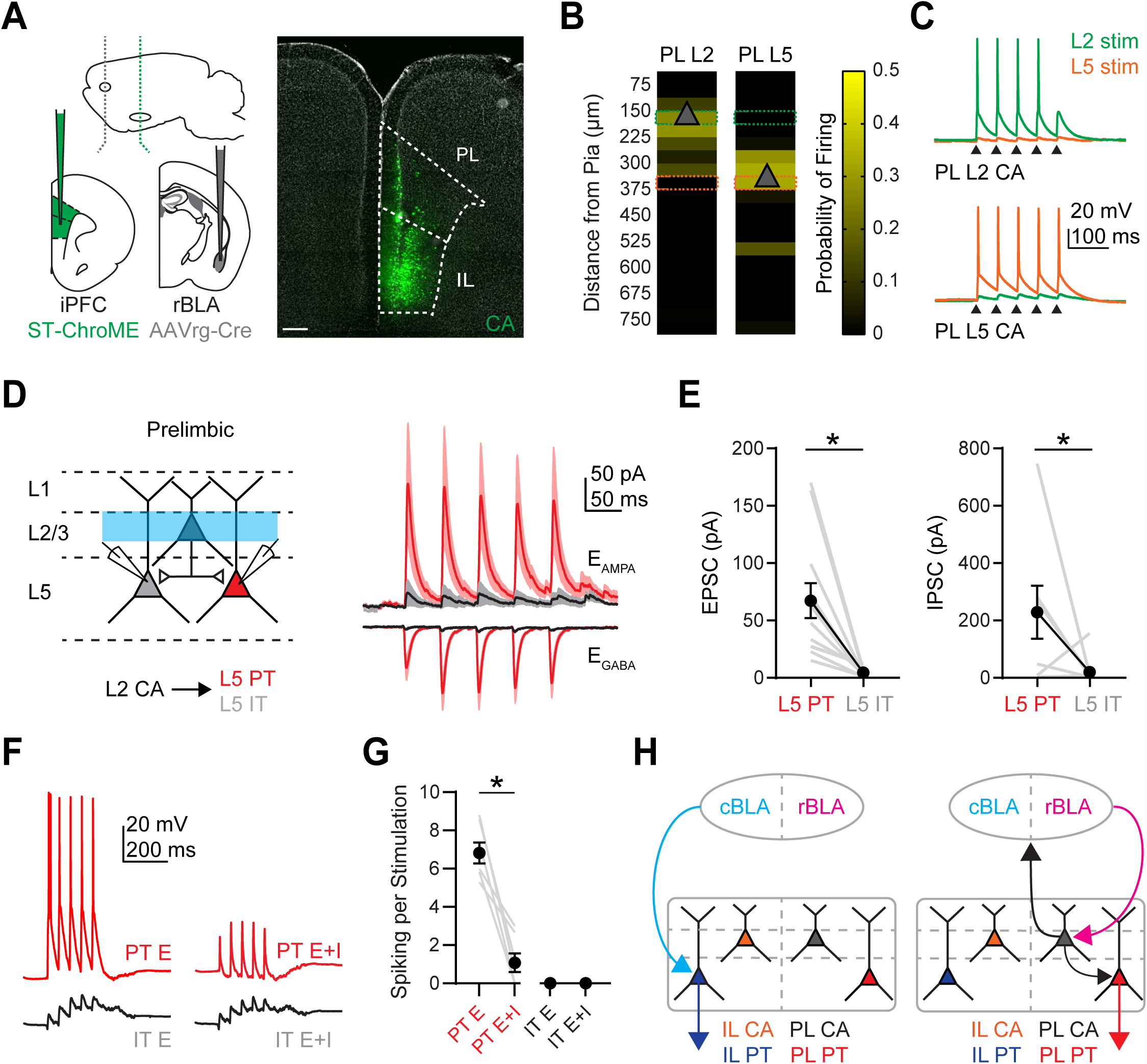
PL L2 CA neurons specifically target PL L5 PT over PL L5 IT neurons. **A)** *Left*, schematic for injections of AAVrg-Cre into rBLA, AAV-DIO-St-ChroME into PFC, CTB-647 into PAG, and CTB-555 into the contralateral PFC. *Right,* Example image of CA neurons retrogradely labeled by AAVrg-Cre and expressing DIO-ChR2-eYFP (green) in the PFC (+2.2 mm from Bregma). Scale bar = 200 µm. DAPI staining is shown in grey. **B)** Laminar stimulation control experiments. Light-evoked St-ChroME-expressing CA neurons in L2 and L5 were recorded in current-clamp and stimulated with 37.5 x 750 µm bars from the pial surface to white matter. Light-evoked firing is plotted as a function of distance from pia for PL L2 CA neurons (left) and PL L5 CA neurons (right). Triangle denotes cell body location. **C)** Example traces from L2 (green) or L5 (orange) stimulation of PL L2 CA neurons (top) and PL L5 CA neurons (bottom). Black triangles denote light stimulation. **D)** *Left,* Experimental scheme for recording PL L2 CA local connections. Pairs of PL L5 IT and PL L5 PT neurons were recorded while stimulating PL L2 CA neurons at 20 Hz. *Right,* Average PL L2 CA-evoked EPSCs and IPSCs recorded at PL L5 IT (black) and PL L5 PT (red) neurons. **E)** *Left,* Summary of PL L2 CA-evoked EPSC_1_ amplitudes at PL L5 IT and PL L5 PT neurons, where grey lines denote individual pairs. *Right,* Similar for IPSC_1_ amplitudes (n = 12 pairs for EPSCs and 10 pairs for IPSCs, 7 animals). **F)** Example traces from dynamic-clamp experiments, where excitatory (E) or mixed excitatory and inhibitory (E+I) conductances derived from (D-E) were injected into the respective L5 PT (red) or IT (black) cells. **G)** Summary of dynamic-clamp experiments at maximum scaled conductance values, showing the average spikes elicited from injecting E or E+I conductances into respective cell-types. Grey lines denote spiking from individual neurons. (n = 7 for each cell-type, 4 animals). **H)** *Left,* Revised model of cBLA to IL synaptic connectivity. *Right*, Revised model of rBLA to PL synaptic connectivity. * p < 0.05

Lastly, we used dynamic-clamp recordings to determine how PL L2 CA activation influence the firing properties of PL L5 PT and IT cells (Carter & Regehr, 2002; McGarry & Carter, 2016; Anastasiades *et al*., 2018). We injected scaled conductances from our voltage-clamp recordings, testing if these inputs can drive spiking at L5 PT and L5 IT cells (**Fig. 6F & S7**). Injecting excitatory conductances alone resulted in robust spiking in PT cells but not IT cells, whereas injecting combined excitatory and inhibitory conductances resulted in significantly attenuated spiking of PT cells (spikes per stimulus: PT E = 6.82 **±** 0.54; PT E + I = 1.07 **±** 0.49; IT E = 0; IT E + I = 0) (**Fig. 6G**). These results indicate that activation of PL L2 CA neurons drives PL L5 PT spiking but this can be regulated by local inhibitory circuits. Together, these results describe the multiple divergent pathways through which rBLA and cBLA influence activity in the PFC, indicating that different sub-regions of the BLA regulate multiple PFC outputs via both mono- and polysynaptic pathways (**Fig. 6H**).

## DISCUSSION

We have determined how connections from rBLA and cBLA engage different subregions, layers, and cell types in the PFC. Distinct amygdalocortical neurons in the rBLA and cBLA send divergent axonal projections to the PL and IL. rBLA activates PL L2, but also generates mixed excitation and inhibition across L5, whereas cBLA primarily activates IL L5. Synaptic connections are cell-type specific, with rBLA primarily contacting and driving PL L2 CA neurons, and cBLA mainly engaging IL L5 PT neurons. rBLA also evokes polysynaptic responses in PL L5 PT neurons via PL L2 CA neurons, allowing these inputs to influence subcortical output. Together, our findings reveal several new levels in the organization of distinct rBLA-PFC and cBLA-PFC circuits.

The BLA is integral for emotional behaviors, with function varying across the rostral-caudal axis (Senn *et al*., 2014; Kim *et al*., 2016; Beyeler *et al*., 2018). We focused on the rostral and caudal poles of the BLA, which roughly correspond to the anatomically segregated and functionally opposed anterior BLA and posterior BLA (Kim *et al*., 2016; Kim *et al*., 2017; Pi *et al*., 2020). In support of this model of BLA organization, we found that BLA to PFC connectivity is strongly divergent to target sub-regions and layers. Interestingly, equivalent connectivity is also found in primates, with magnocellular BLA projecting to superficial layers of area 32, a primate analogue of rBLA projecting to PL, and parvocellular BLA projecting more diffusely to L5 of area 25, a primate analogue of cBLA projecting to IL (Sharma *et al*., 2020). This homologous circuit organization suggests that divergent rBLA and cBLA projections to the PFC is a conserved element of amygdalocortical connectivity. There is likely additional diversity in these circuits, as gradients in cell-types and projection targets are also observed in the medial-lateral and ventral-dorsal axes of the BLA (McGarry & Carter, 2017; Beyeler *et al*., 2018; O’Leary *et al*., 2020).

BLA activity *in vivo* can evoke both excitation and inhibition of PFC neurons in a valence-specific manner (Burgos-Robles *et al*., 2017). In our *in vivo* recordings, we found that stimulation of rBLA and cBLA also causes distinct patterns of activation and suppression across sub-regions and layers of the PFC. rBLA exerts broad influence over most of PFC, with strong activation of PL L2, but also suppression of PL L5 and IL L5. In contrast, cBLA generates a mix of activation and suppression in IL L5, but has only minimal influence on either PL L2 or PL L5. This evoked activity does not reflect a simple gradient of rBLA to dorsal PFC and cBLA to ventral PFC, with the divergence of L5 responses highlighting how rBLA and cBLA are not merely parallel pathways from a single input. IL L5 is particularly important for threat processing (Adhikari *et al*., 2015; Bukalo *et al*., 2015; Bloodgood *et al*., 2018), and we find it can be inhibited by rBLA but excited by cBLA inputs. In the future, it will be particularly interesting to determine how rBLA and cBLA projections to the PFC encode emotional valence, including threat processing.

Our slice physiology revealed cell-type specific connectivity, with rBLA primarily targeting reciprocally projecting PL L2 CA neurons, and cBLA almost exclusively targeting subcortically projecting IL L5 PT neurons that directly influence subcortical nuclei involved in threat responses (Do-Monte *et al*., 2015; Vander Weele *et al*., 2018; Huang *et al*., 2019).. These findings reconcile previous studies, which showed how non-spatially restricted BLA inputs target PL L2 CA neurons (Little & Carter, 2013) and IL L5 PT neurons (Cheriyan *et al*., 2016). While IL L2 CA neurons also receive input from both rBLA and cBLA, these connections are weaker than onto the primary targets. Interestingly, PL L5 PT neurons receive no direct input from either rBLA or cBLA, even though they are strongly implicated in mediating threat detection (Rozeske *et al*., 2018; Huang *et al*., 2019). Instead, we found PL L5 PT neurons receive polysynaptic excitation from rBLA via PL L2 CA neurons, as well as inhibition via the local circuit. These findings provide a mechanism for how differences in cell-type specific targeting and activation from BLA to PL and IL arise from different presynaptic BLA sub-regions.

Several of our experiments indicate that rBLA and cBLA inputs evoke distinct patterns of excitation and inhibition across subregions and layers of the PFC. Our slice recordings show rBLA evokes inhibition in multiple cell types, including PL L2 CA, PL L5 PT and IL L5 PT neurons. However, the E/I ratio was higher for PL L2 CA neurons, consistent with our *in vivo* studies showing this subregion and layer is primarily excited by these inputs. In contrast, our slice recordings show cBLA evokes prominent excitation and inhibition only at IL L5 PT neurons, again consistent with our *in vivo* studies. In the future, it will be important to establish which interneurons mediate rBLA and cBLA-evoked inhibition in L5 across sub-regions of the PFC, and determine if they are similar to cells targeted by glutamatergic inputs from contralateral cortex (Anastasiades *et al*., 2018), thalamus (Anastasiades *et al*., 2021), or hippocampus (Liu *et al*., 2020).

Repetitive activation of rBLA and cBLA inputs generate cell-type specific firing in the PFC, allowing them to activate distinct output pathways either directly or through the local circuit. rBLA drives PL L2 CA neurons, which in turn cause polysynaptic excitation and inhibition in PL L5 PT neurons, consistent with previous work showing BLA engages in a reciprocal loop with PFC (Little & Carter, 2013; McGarry & Carter, 2017). In contrast, cBLA strongly activates IL L5 PT neurons, has minimal influence on PL L2 CA or IL L2 CA neurons, and generates minimal polysynaptic responses in PL L5. Interestingly, this evoked activity is different from ventral hippocampal inputs, which are another major afferent to IL that primarily engage L5 IT neurons (Liu and Carter, 2018). Our finding of minimal inter-subregion connectivity contrasts with previous work showing connections from IL L5/6 to PL L5/6 (Marek *et al*., 2018). One explanation is that communication is mediated by L5 IT cells, which are poorly activated by cBLA inputs (**Fig. S5**). In contrast, L5 PT cells are a major output pathway across the cortex (Economo *et al*., 2018), and have relatively few connections locally (Morishima & Kawaguchi, 2006), consistent with little impact of cBLA activity on PL.

As a mechanism for how signals spread in the PFC, we found that PL L2 CA neurons project to PL L5 PT neurons and not neighboring PL L5 IT neurons. Interestingly, these connections bypass L3, reminiscent of superficial cells crossing L4 when projecting to deeper layers of sensory cortex (Lefort *et al*., 2009; Hooks *et al*., 2011). The extreme bias in targeting of L5 PT over L5 IT also agrees with targeted connectivity observed in other parts of frontal cortex (Anderson *et al*., 2010). It also suggests a fundamental difference in L2/3 connectivity within the local circuit between sensory and frontal cortices (Hirai *et al*., 2012; Joshi *et al*., 2015). Together, our findings indicate that PL L2 CA neurons serve as a crucial node in a disynaptic feedforward circuit, which links rBLA input to PL L5 PT cells. Consequently, the influence of rBLA on PFC output will depend on the neuromodulatory state, which can alter the tightly balanced excitation and inhibition onto PL L5 PT cells (Floresco & Tse, 2007; Vander Weele *et al*., 2018; Anastasiades *et al*., 2019).

Long-range circuits exist in multiple motifs, including unidirectional connections (Liu & Carter, 2018), “closed” reciprocal loops (Little & Carter, 2013), and “open” reciprocal loops (Collins *et al*., 2018). Our results confirm that rBLA participates in a rare example of a closed reciprocal loop with PL, contacting L2 CA neurons that project back to the rBLA (Little and Carter, 2013). These connections are under inhibitory control via local interneurons, which prevent recurrent excitation (McGarry and Carter, 2016). In contrast, cBLA does not participate in a reciprocal loop with IL, instead making unidirectional connections onto L5 PT neurons. These connections differ from vHPC inputs that are also unidirectional but instead contact IL L5 IT neurons (Liu & Carter, 2018), as well as thalamic inputs that instead contact L2/3 pyramidal cells (Collins *et al*., 2018) and the apical dendrites of L5 PT neurons (Anastasiades *et al*., 2021). Instead, cBLA inputs appear most similar to callosal inputs from the contralateral PFC, which span multiple layers, but within L5 are also biased onto PT neurons, although not to the same degree (Anastasiades *et al*., 2018; Anastasiades & Carter, 2021).

In conclusion, our results determine how rBLA and cBLA make distinct projections to PL and IL, driving the activity of different output pathways via two unique circuits. Along with recent studies (Kim *et al*., 2016; Zhang *et al*., 2021), our results contribute to a new framework to study the role of BLA-PFC circuits, with implications for behavior. For example, BLA to PL projections are necessary for expression of threat conditioning (Senn *et al*., 2014; Burgos-Robles *et al*., 2017), and because rBLA is the main input to PL, our findings are consistent with the rBLA playing a key role in aversion (Kim *et al*., 2016; Pi *et al*., 2020). In contrast, BLA to IL projections are involved in the extinction of threat conditioning (Senn *et al*., 2014; Klavir *et al*., 2017), and because cBLA is the primary input to IL, our findings are also consistent with cBLA being involved in this learning (Zhang *et al*., 2020). In the future, it will be important to explicitly test these ideas by measuring activity in specific sub-circuits of the BLA and PFC during different forms of motivated behavior.

## MATERIALS & METHODS

Experiments involved P28-P70 wild-type mice on a C57 BL/6J background. Experiments used male and female mice, and no significant differences were found between groups. All procedures followed guidelines approved by the New York University Animal Welfare Committee.

### Stereotaxic injections

P28-P56 mice were deeply anesthetized with either isoflurane or a mixture of ketamine and xylazine and then head fixed in a stereotax (Kopf Instruments). A small craniotomy was made over the injection site, through which retrograde tracers and/or viruses were injected using a Nanoject III (Drummond). Injection site coordinates were relative to bregma (mediolateral, dorsoventral, and rostrocaudal axes: Prelimbic PFC = ±0.35, -2.1, +2.2 mm; Infralimbic PFC = ±0.35, -2.3, +2.2 mm; rBLA = -3.0, -5.1, -1.1 mm; cBLA = -3.0, -5.1, -1.7 mm; PAG = -0.6, -3.0 and -2.5, -4.0 mm). Dual AAVrg experiments were injected at +2.4 mm for PL and +2.0 mm for IL in the rostrocaudal axis to minimize viral leak. Borosilicate pipettes with 5-10 µm tip diameters were back-filled, and 100-500 nL was pressure-injected, with 30-45 second inter-injection intervals.

For retrograde labeling or confirmation of injection site for electrophysiology experiments, pipettes were filled with either cholera toxin subunit B (CTB) conjugated to Alexa 488, 555, or 647 (Life Technologies) or blue latex beads. AAV1-hSyn-hChR2-eYFP (UPenn Vector Core AV-1-26973P / Addgene 26973-AAV1) or AAV1-CamKIIa-hChR2-mCherry (UPenn Vector Core AV-1-26975 / Addgene 26975-AAV1) were used for non-conditional axon labeling. AAVrg-CAG-tdTomato (Addgene 59462-AAVrg) and AAVrg-CAG-GFP (Addgene 37825-AAVrg) were used for retrograde labeling for histological experiments. Optogenetic stimulation was achieved using AAV1-hSyn-hChR2-eYFP, AAV1-EF1a-DIO-hChR2-eYFP (UPenn Vector Core AV-1-20298P / Addgene 20298-AAV1) or AAV9-CAG-DIO-ChroME-ST-p2A-H2B-mRuby (generously provided by Hillel Adesnik). The St-ChroME virus was diluted 1:10 in 0.01M PBS prior to injection. Simultaneous virus and tracer injections were mixed in a 1:1 virus:tracer ratio, except for experiments involving axonal stimulation evoked firing, which used a 3:2 virus:tracer ratio. Following injections, the pipette was left in place for an additional 10 min before being slowly withdrawn to ensure injections remained local. Retrograde-Cre experiments to label cortico-amygdalar (CA) neurons were carried out in a similar manner, with injection of AAVrg-hSyn-Cre (Addgene 105553-AAVrg). After all injections, animals were returned to their home cages for 2-4 weeks before being used for experiments, except for experiments involving st-ChroME virus, which were returned for 7-9 days to minimize expression time.

### In vivo electrophysiology

Two to four weeks after viral injections into either rBLA or cBLA, mice were anesthetized with isoflurane, the skin overlying the skull was surgically removed, and a custom plate was attached to the skull using Meta-bond (Parkell), leaving the area over the PFC exposed. An AgCl reference electrode was implanted contralateral to the recording site, attached to an external gold pin. A multimode, 200 µm ID fiber optic connected to a 1.25 mm steel ferrule (Thorlabs) was implanted in either the rBLA or cBLA. The exposed skull was then covered with Kwik-Cast (WPI), and mice were allowed to recover in the home cage for at least 3 days before recording.

On the day of the recording, mice were anesthetized with isoflurane and a small craniotomy was made over the PFC, using the stereotaxic coordinates designated above. After recovering from anesthesia, mice were then head-fixed using the implanted plate and allowed to run freely on a spinning wheel treadmill. A 473 nm LED was attached to the ferrule with a 200 µm ID patch cable (Thorlabs), allowing stimulation of rBLA or cBLA. Before insertion into the brain, a Neuropixels electrode array (Jun *et al*., 2017) was mounted on a micromanipulator (Sutter Instruments) and painted with DiI (Thermofisher, 2 mg/mL in ethanol) for post-hoc track reconstruction. The Neuropixels probe was lowered vertically 3 mm below the dorsal brain surface and, laterally from the midline, either 350-450 µm for L5 recordings or 150-250 µm for L2/3 recordings. The probe was advanced slowly (∼2 µm/s) and allowed to rest for 30 minutes before recording. A drop of silicone oil (1000cs, Dow Corning) was placed over the exposed brain to prevent drying. Probes were referenced to the implanted Ag/AgCl wire. Analog traces were filtered (0.3 – 5 kHz), digitized, and recorded (30 kHz per channel) using acquisition boards from National Instruments and OpenEphys software (Siegle *et al*., 2017). Neuropixels recordings were made in external reference mode with LFP gain of 250 and AP gain of 500. Either rBLA or cBLA was photo-stimulated at 20 Hz for 5 pulses of 2 ms duration, repeated for 40 trials with an inter-trial interval of 30 seconds. Following recordings, brains were collected and processed for post-hoc probe track reconstruction.

### Slice preparation

Mice were anesthetized with an intraperitoneal injection of a lethal dose of ketamine and xylazine and then perfused intracardially with an ice-cold cutting solution containing the following (in mM): 65 sucrose, 76 NaCl, 25 NaHCO_3_, 1.4 NaH_2_PO_4_, 25 glucose, 2.5 KCl, 7 MgCl_2_, 0.4 Na-ascorbate, and 2 Na-pyruvate (bubbled with 95% O_2_/5% CO_2_). 300 µm coronal sections were cut in this solution and transferred to ACSF containing the following (in mM): 120 NaCl, 25 NaHCO_3_, 1.4 NaH_2_PO_4_, 21 glucose, 2.5 KCl, 2 CaCl_2_, 1 MgCl_2_, 0.4 Na-ascorbate, and 2 Na-pyruvate (bubbled with 95% O_2_/5% CO_2_). Slices were recovered for 30 min at 35°C and stored for at least 30 min at 24°C. All experiments were conducted at 30-32°C.

### Slice electrophysiology

Targeted whole-cell recordings were made from projection neurons in PL and IL using infrared-differential interference contrast. In the PFC, layers were defined by distance from the pial surface: L2: 170-220 µm; L3: 225-300 µm; L5: 350-550 µm; L6: 600-850 µm. PL and IL were defined visually, and the intermediate ventral PL area was deliberately avoided. CA, CC, and PT neurons were identified by the presence of fluorescently tagged CTB. For recordings from st-ChroME+ CA neurons, cells in L2 or L5 were identified by mRuby expression.

For voltage-clamp experiments, borosilicate pipettes (3-5 MΩ) were filled with (in mM): 135 Cs-gluconate, 10 HEPES, 10 Na-phosphocreatine, 4 Mg_2_-ATP, 0.4 NaGTP, 10 CsCl, and 10 EGTA, pH 7.3 with CsOH (290-295 mOsm). For current-clamp and dynamic-clamp recordings, borosilicate pipettes (3-5 MΩ) were filled with (in mM): 135 K-gluconate, 7 KCl, 10 HEPES, 10 Na-phosphocreatine, 4 Mg_2_-ATP, 0.4 NaGTP, and 0.5 EGTA, pH 7.3 with KOH (290-295 mOsm). In some voltage-clamp experiments, 1 µM TTX was included in the bath to block action potentials (APs), along with 0.1 mM 4-AP and 4 mM external Ca^2+^ to restore presynaptic glutamate release. In some current-clamp experiments, 10 µM CPP and 10 µM gabazine were included to block NMDA receptors and GABA_A_ receptors, respectively. In voltage-clamp experiments recording IPSCs, 10 µM CPP was included to block NMDA receptors. In dynamic-clamp recordings, 10 µM NBQX, 10 µM CPP, and 10 µM gabazine were included to block AMPA, NMDA, and GABA_A_ receptors, respectively. All chemicals were from Sigma or Tocris Bioscience.

Electrophysiology data for voltage-clamp and current-clamp experiments were collected with a Multiclamp 700B amplifier (Axon Instruments) and National Instruments boards using custom software in MATLAB (MathWorks). Dynamic-clamp recordings were performed using an ITC-18 interface (Heka Electronics) running at 50 kHz with Igor Pro software (Wavemetrics) running MafPC (courtesy of Matthew Xu-Friedman). For dynamic-clamp recordings, conductances were initially converted from st-ChroME local-circuit mapping experiments and then injected into neurons while being multiplied by a range of constant scale-factors to model the effects of driving the L2 CA input at various stimulus strengths. The reversal potentials for AMPA-R excitation and GABAa-R inhibition were set to 0 mV and -75 mV, respectively. Signals were sampled at 10 kHz and filtered at either 5 kHz for current-clamp and dynamic-clamp recordings, or 2 kHz for voltage-clamp recordings. Series resistance was measured as 10-25 MΩ and not compensated.

### Slice optogenetics

Glutamate release was triggered by activating channelrhodopsin-2 (ChR2) present in the presynaptic terminals of rBLA or cBLA inputs to the PFC (Little & Carter, 2012; McGarry & Carter, 2016). LED power was adjusted until responses >100 pA were seen in at least one neuron in a recorded pair or triplet, with the same power used for all cells in that slice. ChR2 was activated with 2 ms pulses of 473 nm light from a blue light-emitting diode (LED; 473 nm; Thorlabs) through a 10X 0.3 NA objective (Olympus) with a power range of 0.4-12 mW. In some experiments, ChR2 was activated by 20 Hz stimulation with 2 ms pulses of light. Subcellular targeting recordings utilized a digital mirror device (Mightex Polygon 400G) to stimulate a 10x10 grid of 75-micron squares at a power range of 0.05-0.2 mW per square at 1 Hz with the first row aligned to the pia. For all other recordings, the objective was centered over the soma unless noted otherwise. Inter-trial interval was 10 seconds except for experiments involving trains of stimulation, in which it was 30 seconds.

### Soma-restricted optogenetics

To map the outputs of st-ChroME+ CA neurons, stimulation parameters were first developed to produced robust, spatially restricted AP firing. Recordings were made from PL L2 and L5 st-ChroME+ CA neurons located in the same slice of PFC. Blue (473 nm) LED light was illuminated as 37.5 x 750 µm bars using a DMD through a 10X 0.3 NA objective. Individual bars were stimulated with 5 pulses of 1 ms light at 20 Hz with an intensity of 0.1-0.8 mW in a pseudorandom order. Responses were recorded at resting membrane potentials. High power (0.8 mW) stimulation resulted in st-ChroME+ neurons firing outside of their layer and therefore was not used in subsequent experiments. In experiments involving the postsynaptic targeting of CA neurons onto CC and PT neurons, the 4^th^ or 9^th^ bar from the pia, which is equivalent to L2/3 or L5, were alternatingly stimulated at 0.1-0.4 mW, with a 15 second inter-stimulus interval.

### Histology

Mice were anesthetized and perfused intracardially with 0.01 M PBS followed by 4% PFA. Brains were stored in 4% PFA for 12-18 hours at 4°C before being washed three times (30 minutes each) in 0.01 M PBS. Slices were cut on a VT-1000S vibratome (Leica) at 100 µm thickness, except for Neuropixels tract reconstruction which were cut at 70 µm thickness, and placed on gel-coated glass slides. ProLong Gold anti-fade reagent with DAPI (Invitrogen) or VectaShield with DAPI (Vector Labs) was applied to the surface of the slices, which were then covered with a glass coverslip. Fluorescent images were taken on an Olympus VS120 microscope, using a 10X 0.25 NA objective (Olympus). For rBLA and cBLA axonal projections to the PFC and retrograde cell counting in the BLA, images were taken on a TCS SP8 confocal microscope (Leica), using a 10X 0.4 NA objective or 20x 0.75 NA objective (Leica).

### Data analysis

Analysis of slice electrophysiology was performed using Igor Pro (WaveMetrics), except for MATLAB (MathWorks) for st-ChroME control experiments and sCrACM recordings. For voltage-clamp recordings, PSC amplitudes were measured as the average at 1 ms around the peak response.

*In vivo* electrophysiology data were pre-processed by referencing to the common median across all channels. The data were then spike-sorted using Kilosort2 (Pachitariu *et al*., 2016), (https://github.com/MouseLand/Kilosort2), with manual curation in Phy (https://github.com/cortex-lab/phy). Spike time and waveform data were further processed and visualized using MATLAB code modified from N. Steinmetz (https://github.com/cortex-lab/spikes). Importantly, all units were then aligned to brain location using the Allen common coordinate framework (CCF) (Wang *et al*., 2020). Significantly responsive single units were determined by using a Wilcoxon signed-rank test (threshold of p < 0.05) comparing the number of spikes in a 2 second baseline period before LED stimulation and the number of spikes 5 to 25 ms after every LED pulse. If the unit had significantly lower firing during the LED window and an average z-scored value of -0.2 or less, it was designated as inhibited, and if the unit had significantly higher firing during the LED and an average z-scored value of +0.2 or more, it was designated as significantly excited. Binomial tests were used to compare the proportion of significantly excited or inhibited units across PFC in response to rBLA or cBLA stimulation and were corrected for multiple comparisons by the Holm-Bonferroni method.

Summary data are reported in the text and figures as arithmetic mean ± SEM or geometric mean and geometric SD factor for ratio data. Ratio data displayed in figures on logarithmic axes are reported as geometric mean ± geometric SD. In some graphs with three or more traces, SEM waves are omitted for clarity. Statistical analyses were performed using Prism 9 (GraphPad Software). Comparisons between unpaired data were performed using two-tailed Mann-Whitney tests. Comparisons between data recorded in pairs were performed using two-tailed Wilcoxon matched-pairs signed rank tests. Ratio data were log-transformed and compared to a theoretical median of 0. For paired comparisons of more than two groups, Friedman tests with Dunn’s multiple comparisons tests were performed. For unpaired comparisons of more than two groups, Kruskal-Wallis tests with Dunn’s multiple comparisons tests were performed. For all tests, significance was defined as p < 0.05.

Cell counting in the BLA was performed using ImageJ on a multi-color image of retrogradely labeled neurons. ROIs for rBLA and cBLA were calculated in each slice after aligning to the Allen Brain Atlas at the appropriate rostrocaudal co-ordinate. The number of cells per slice was averaged across 3 animals and used to calculate averages ± SEM across animals. Axon distributions in the PFC were quantified using un-binned fluorescence profiles relative to distance from the pia for 3 slices from each animal. The average fluorescence profile for each animal was peak-normalized before calculating a final average ± SEM across animals.

## Acknowledgements

We thank the Carter lab, Christine Constantinople, and Robert Froemke for helpful discussions and comments on the manuscript. This work was supported by NIH T32 MH019524 (KM) and NIH R01 MH085974 (AGC). The authors have no financial conflicts of interest.

---

**Figure S1:**
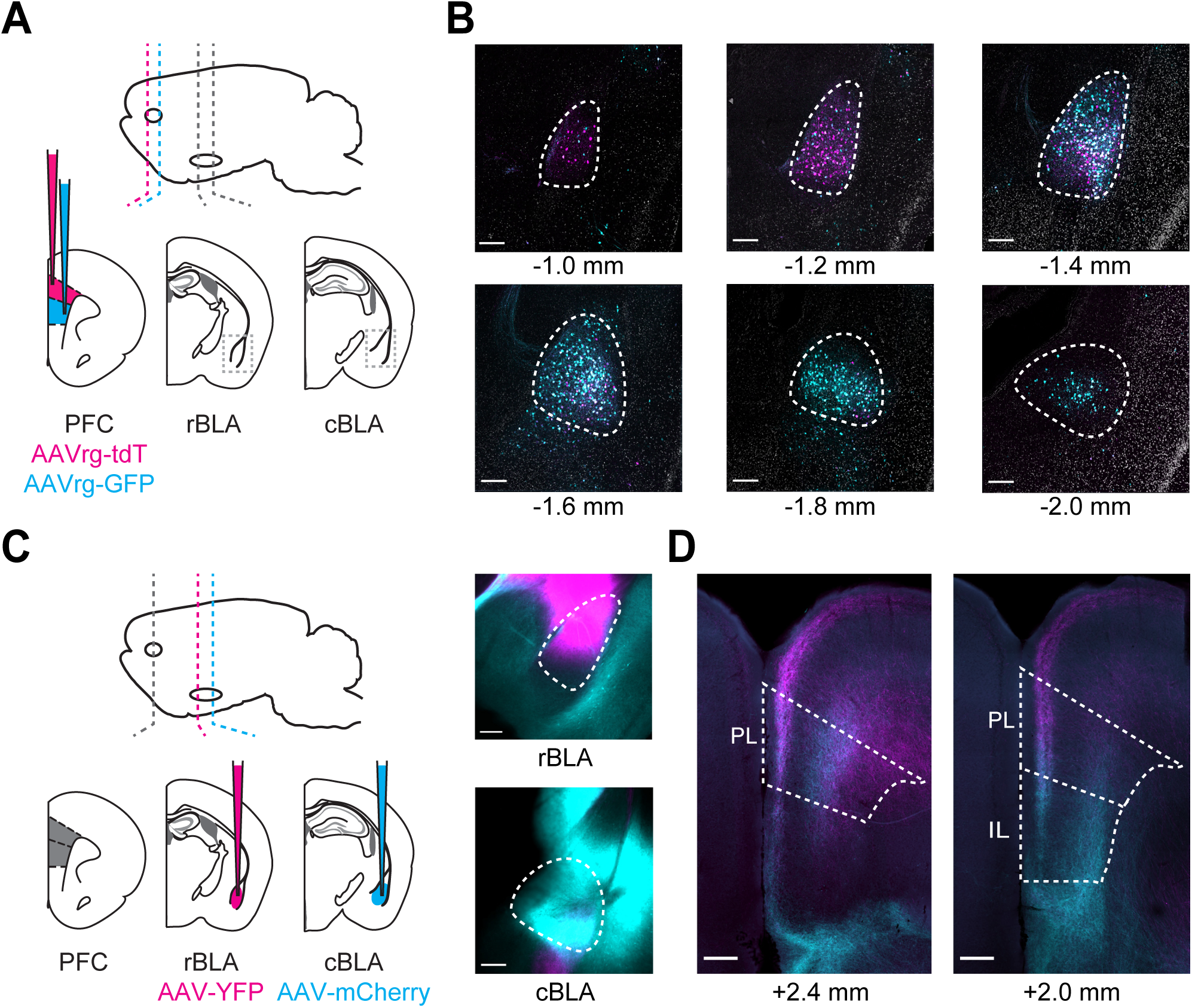
PL and IL-projecting neurons across the BLA. ***Related to Figure 1*** **A)** Schematic for injections of AAVrg-tdTomato into PL (magenta) and AAVrg-GFP into IL (cyan). **B)** Series of representative confocal images of PL-projecting (magenta), IL-projecting (cyan), and DAPI (grey) fluorescent signals in coronal slices across the rostral to caudal extent of BLA. Scale bar = 200 µm. Distance from bregma is shown under the corresponding image. **C)** *Left*, Schematic for injections of AAV-YFP into rBLA (magenta) and AAV-mCherry into cBLA (cyan). *Right*, images of rBLA and cBLA injection sites. Scale bar = 250 µm. **D)** Representative images of rBLA (magenta) and cBLA (cyan) axonal projections to mPFC in coronal slices across the rostral to caudal axis. Scale bar = 250 µm. Distance from bregma is shown under the corresponding image.

**Figure S2.**
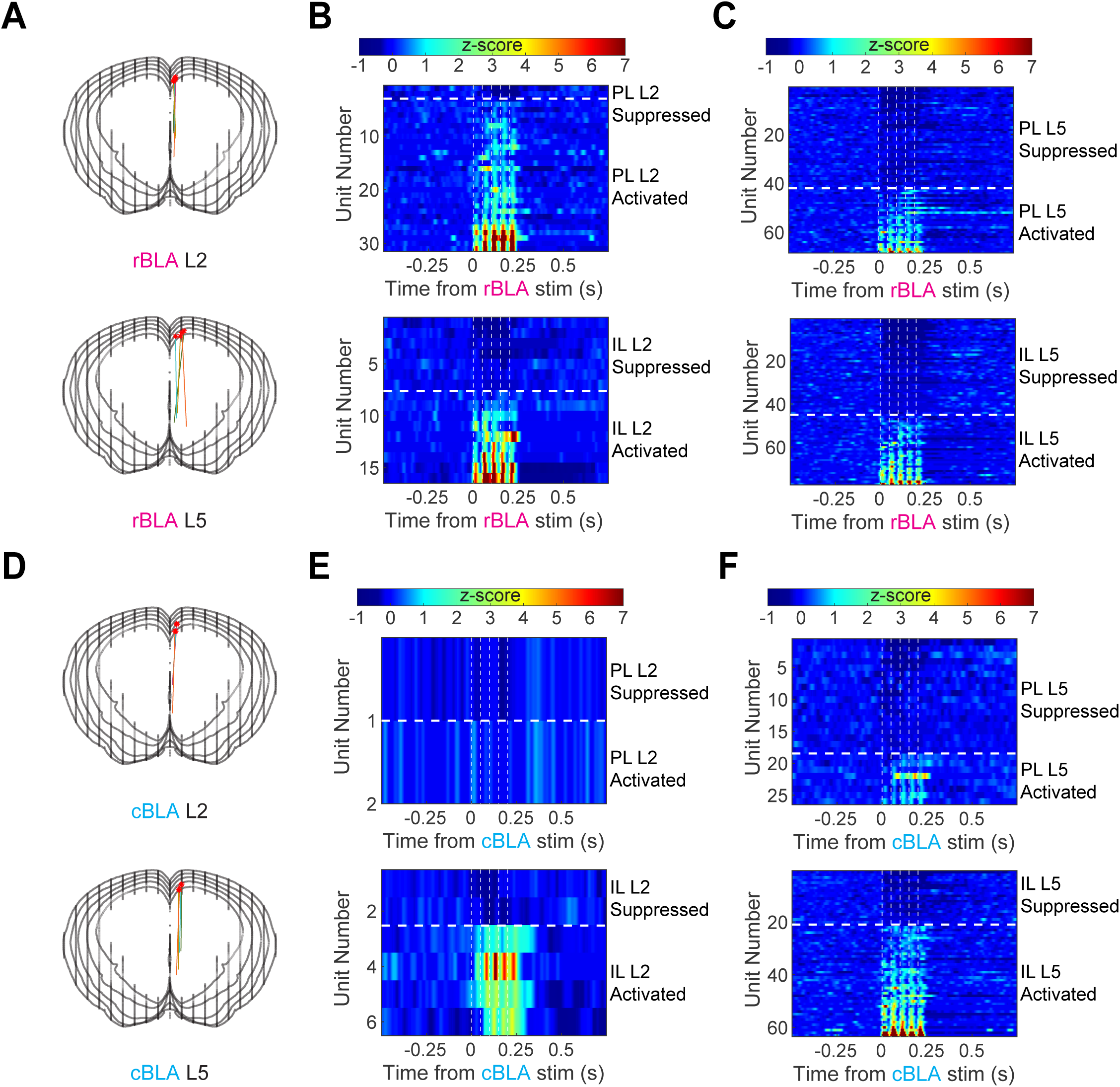
rBLA and cBLA inputs cause distinct patterns of activity across PFC. ***Related to Figure 2*** **A)** *Top,* Reconstructed probe tracts for all L2 insertions from mice injected with ChR2 in rBLA. *Bottom,* Reconstructed probe tracts for all L5 insertions from mice injected with ChR2 in rBLA. **B)** Graph of average responses from each significantly activated or suppressed unit in PL L2 (top) and IL L2 (bottom) in response to rBLA stimulation. **C)** Similar to (B) showing PL L5 and IL L5 responses to rBLA stimulation. **D)** Similar to (A) showing L2 and L5 probe tracts in mice injected with ChR2 into cBLA. **E)** Similar to (B) showing PL L2 and IL L2 responses to cBLA stimulation. **F)** Similar to (B) showing PL L5 and IL L5 responses to cBLA stimulation.

**Figure S3.**
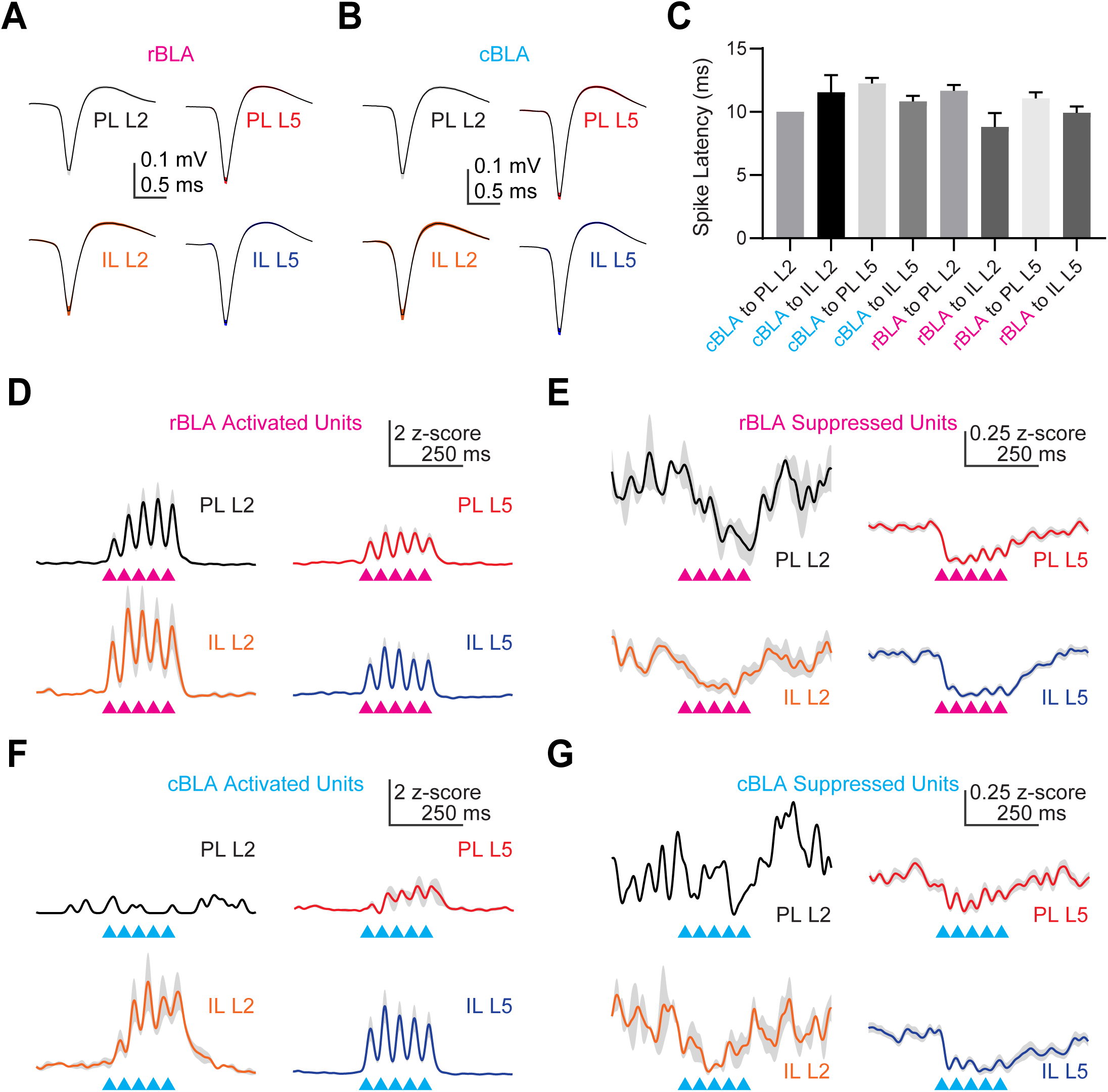
rBLA and cBLA excited and inhibited units. ***Related to Figure 2*** **A)** Average waveform of all single units recorded in PL L2, IL L2, PL L5, and IL L5 from animals with rBLA stimulation. **B)** Similar to (A) for single units recorded from animals with cBLA stimulation. **C)** Graph of average time to spike for significantly activated single units after rBLA or cBLA stimulation within a window of 5-25 ms after every pulse. **D)** Average smoothed, z-scored response from rBLA-activated units in PL L2 (black), IL L2 (orange), PL L5 (red), and IL L5 (blue). **E)** Similar to (D) for rBLA-suppressed units. **F)** Similar to (D) for cBLA-activated units. **G)** Similar to (D) for cBLA-suppressed units.

**Figure S4.**
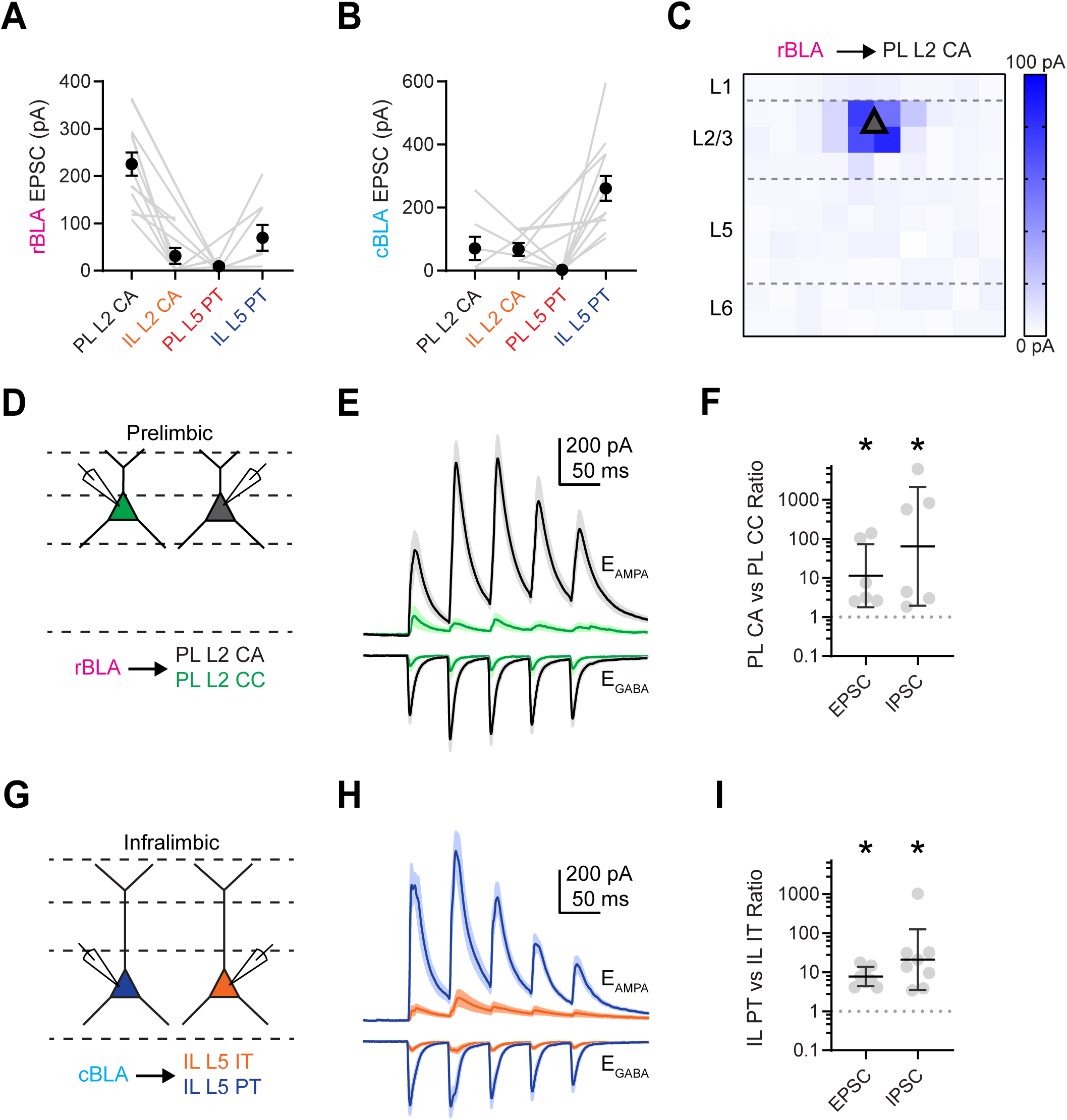
rBLA and cBLA monosynaptic inputs. ***Related to Figure 3*** **A)** Graph of EPSC amplitudes from rBLA to PL L2 CA, IL L2 CA, PL L5 PT, and IL L5 PT recordings. Individual grey lines denote cells recorded in a pair, triplet, or quadruplet. **B)** Similar to (A) showing EPSC amplitudes from cBLA recordings. **C)** Average rBLA-evoked sCrACM EPSC maps for PL L2 CA neurons (5 cells, 3 animals). **D)** Experimental scheme for examining rBLA-evoked cell-type specific connectivity. Pairs of PL L2 CA (black) and PL L2 CC (green) neurons were recorded in voltage clamp at E_GABA_ and E_AMPA_ to measure rBLA-evoked EPSCs and IPSCs while stimulating at 20 Hz. **E)** Average rBLA-evoked EPSCs and IPSCs recorded at E_GABA_ and E_AMPA_ from PL L2 CA (black) and PL L2 CC (green) neurons. **F)** Summary of EPSC_1_ and IPSC_1_ amplitude ratios of PL L2 CA and PL L2 CC neurons from rBLA stimulation. Grey points denote individual pairs. (n = 6 pairs, 4 animals). **G)** Similar to (D) examining cBLA-evoked cell-type specific connectivity in IL L5 focusing on IL L5 PT (blue) and IL L5 IT (orange). **H)** Similar to (E) showing average cBLA-evoked EPSCs and IPSCs from IL L5 PT (blue) and IL L5 IT (orange) neurons. **I)** Similar to (F) showing summary of cBLA-evoked EPSC and IPSC amplitude (n = 8 pairs, 5 animals). * p < 0.05

**Figure S5.**
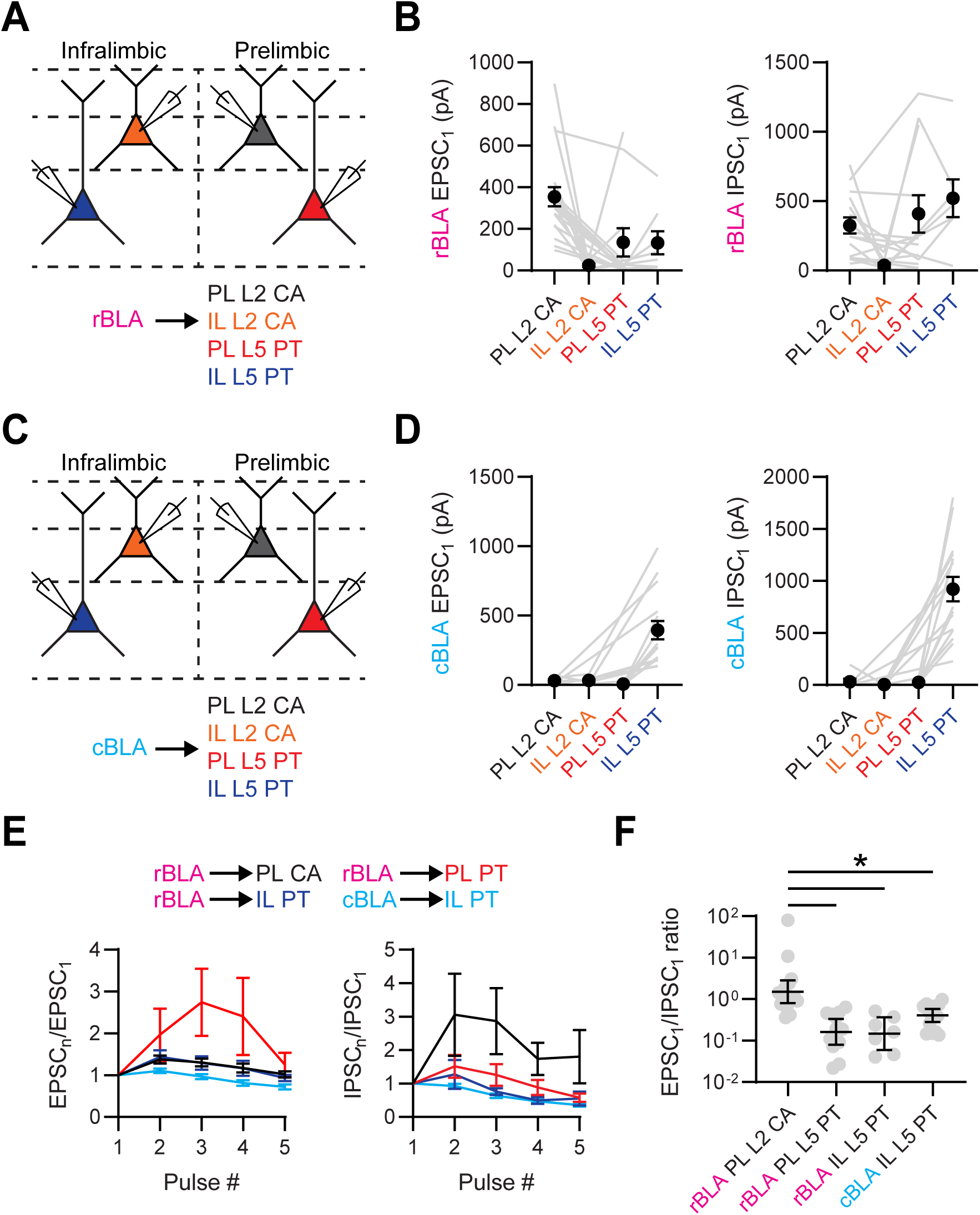
Short-term dynamics of rBLA and cBLA inputs. ***Related to Figure 4*** **A)** Experimental scheme for recording rBLA-evoked short-term dynamics. PL L2 CA (black), IL L2 CA (orange), PL L5 PT (red) and IL L5 PT (blue) neurons. **B)** *Left,* Summary of 1^st^ pulse EPSC amplitudes for rBLA-evoked currents. Grey lines denote individual pairs or triplets. *Right,* similar for 1^st^ pulse IPSC amplitudes for rBLA-evoked currents. **C)** Similar to (A) for recording cBLA-evoked short-term dynamics. **D)** Similar to (B) for cBLA-evoked EPSC and IPSC amplitudes. **E)** *Left,* Summary of EPSC PPR for rBLA-evoked responses onto PL L2 CA (black), PL L5 PT (red), and IL L5 PT (dark blue) neurons as well as cBLA-evoked responses onto IL L5 PT neurons (light blue). *Right,* similar for IPSC PPR. **F)** Summary of E/I ratio (1^st^ pulse EPSC divided by 1^st^ pulse IPSC) for rBLA-evoked responses onto PL L2 CA, PL L5 PT, and IL L5 PT neurons as well as cBLA-evoked responses onto IL L5 PT neurons. Grey points denote individual cells. * p < 0.05

**Figure S6.**
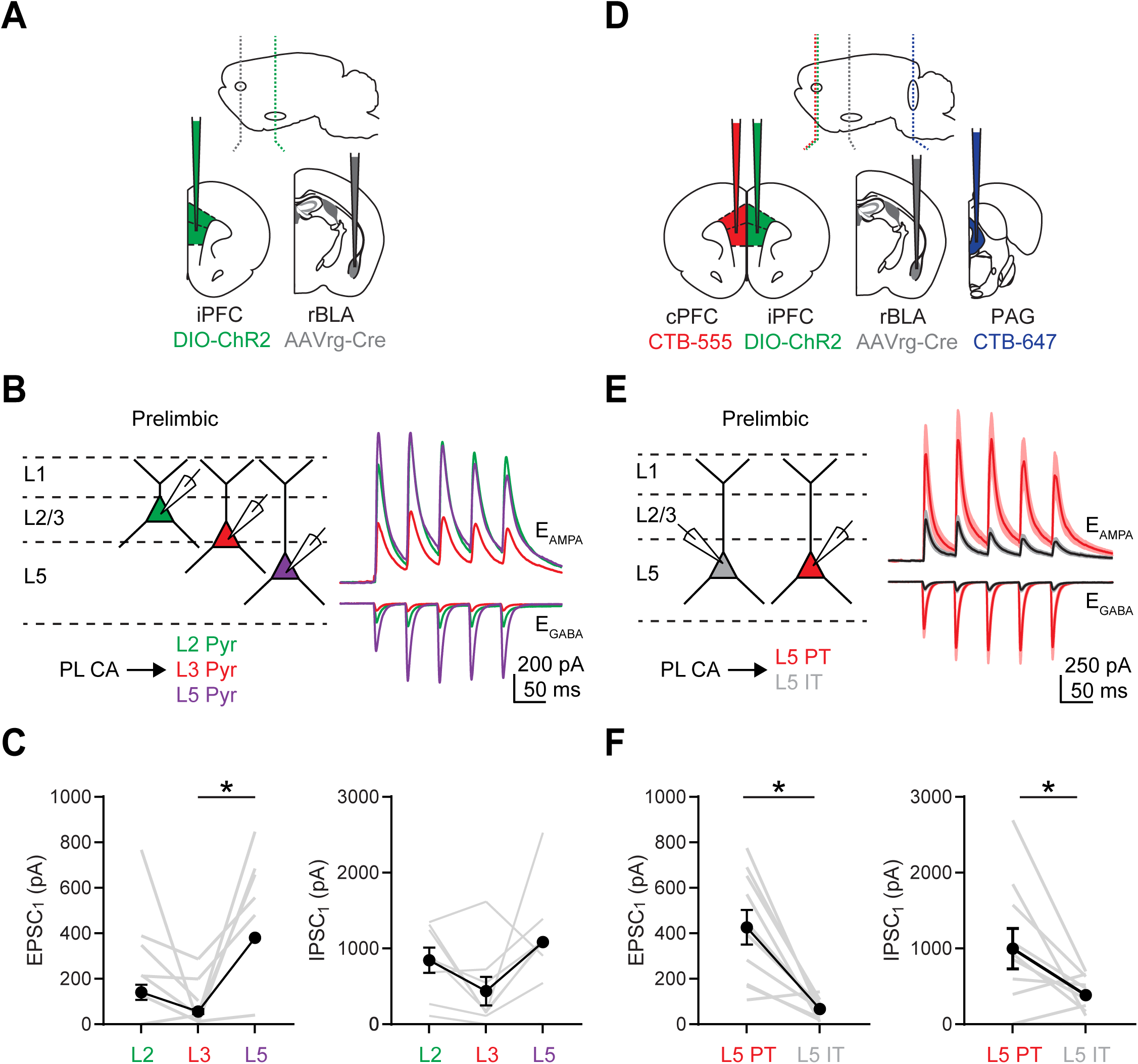
PL CA neurons project within the local network to PL L5 PT neurons. ***Related to Figure 6*** **A)** Schematic for injections of AAVrg-Cre into rBLA and AAV-DIO-ChR2 into ipsilateral PFC. **B)** *Left,* Experimental scheme for recording local CA connectivity. Triplets of L2 (green), L3 (red), and L5 (purple) pyramidal cells were recorded in voltage clamp while stimulating CA neurons at 20 Hz. *Right,* Average CA-evoked EPSCs and IPSCs at PL L2 (green), L3 (red), and L5 (purple) pyramidal cells. **C)** *Left,* Summary of PL CA-evoked EPSC_1_ amplitudes for PL L2, L3, and L5 pyramidal cells, where grey lines denote individual triplets. *Right,* Similar for IPSC_1_ amplitudes (n = 8 triplets, 4 animals). **D)** Schematic for injections of AAVrg-Cre into rBLA, AAV-DIO-ChR2 into PFC, CTB-647 into PAG, and CTB-555 into the contralateral PFC. **E)** *Left,* Experimental scheme for recording local CA connections onto specific L5 neurons. Pairs of PL L5 IT and PL L5 PT neurons were recorded while stimulating CA neurons at 20 Hz. *Right,* Average CA-evoked EPSCs and IPSCs at PL L5 IT (black) and PL L5 PT (red) neurons. **F)** *Left,* Summary of PL CA-evoked EPSC_1_ amplitudes at PL L5 IT and PL L5 PT neurons, where grey lines denote individual pairs. *Left,* Similar for IPSC_1_ amplitudes (n = 10 pairs, 4 animals). * p < 0.05

**Figure S7.**
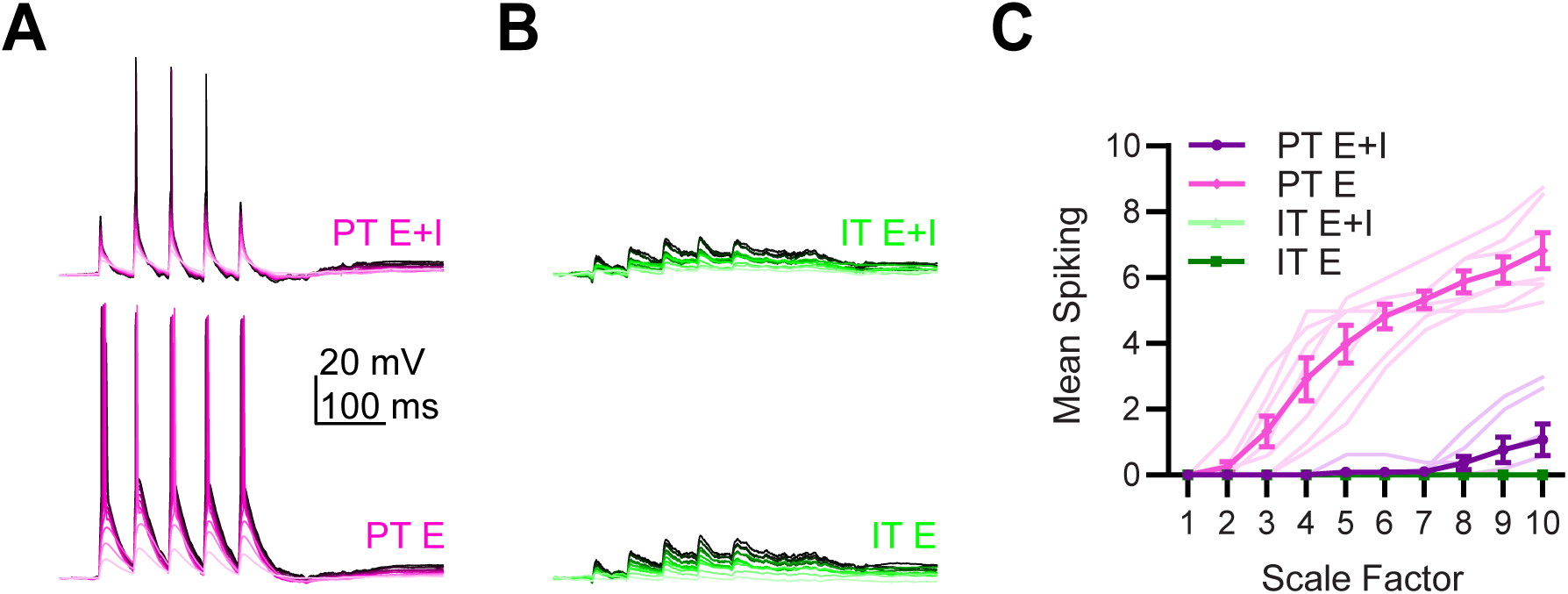
Dynamic-clamp recordings. ***Related to Figure 6*** **A)** Example traces in response to injecting excitatory (E) or mixed excitatory and inhibitory (E+I) conductances into PL L5 PT cells derived from st-ChroME local circuit voltage-clamp experiments. Traces range from light magenta to black based on scale factor (1-10x). **B)** Similar to (A) for PL L5 IT cells (light green to black for each scale factor). **C)** Summary of average dynamic-clamp induced spiking from E and E+I conductances injected into PL L5 PT and IT cells. * p < 0.05

